# Genetic and metabolite biomarkers reveal actinobacteria-mediated estrogen biodegradation in urban estuarine sediment

**DOI:** 10.1101/2020.10.07.329094

**Authors:** Tsun-Hsien Hsiao, Yi-Lung Chen, Menghsiao Meng, Meng-Rong Chuang, Masae Horinouchi, Toshiaki Hayashi, Po-Hsiang Wang, Yin-Ru Chiang

**Affiliations:** Biodiversity Research Center, Academia Sinica, Taipei 115, Taiwan; Department of Microbiology, Soochow University, Taipei 111, Taiwan; Graduate Institute of Biotechnology, National Chung Hsing University, Taichung 402, Taiwan; Condensed Molecular Materials Laboratory, RIKEN, Saitama, 351-0198 Japan; Environmental Molecular Biology Laboratory, RIKEN, Saitama, 351-0198 Japan; Gradaute Institute of Environmental Engineering, National Central University, Taoyuan 320, Taiwan; Earth-Life Science Institute (ELSI), Tokyo Institute of Technology, Tokyo, Japan

**Keywords:** actinobacteria, biodegradation, biomarkers, estrogen biodegradation, oxygenases, *Rhodococcus*, steroid hormones, estuary bioremediation

## Abstract

Steroidal estrogens are often accumulated in urban estuarine sediments worldwide at microgram per gram levels. These aromatic steroids have been classified as endocrine disruptors with an EC50 at sub-nanomolar concentrations and classified as Group 1 carcinogens by the World Health Organization. Microbial degradation is a naturally occurring mechanism that mineralizes estrogens in the biosphere; however, the corresponding genes in estrogen-degrading actinobacteria remain unidentified. In this study, we identified a gene cluster encoding several putative estrogen-degrading genes in actinobacterium *Rhodococcus* sp. strain B50. Among them, the *oecB* and *oecC* genes involved in estrogenic A-ring cleavage were identified through gene-disruption experiments. We also detected the accumulation of two extracellular estrogenic metabolites, including pyridinestrone acid (PEA) and 3aα-H-4α(3’-propanoate)-7aβ-methylhexahydro-1,5-indanedione (HIP), in the estrone-fed strain B50 cultures. Since actinobacterial *oecC* and proteobacterial *oecC* shared less than 40% sequence identity, *oecC* could serve as a specific biomarker to differentiate the contribution of actinobacteria and proteobacteria in environmental estrogen degradation. Therefore, *oecC* and the extracellular metabolites PEA and HIP were used as biomarkers to investigate estrogen biodegradation in an urban estuarine sediment. Interestingly, our data suggested that actinobacteria, rather than alpha-proteobacteria function in sewage treatment plants, are actively degrading estrogens in the urban estuarine sediment.

**Graphical Abstract:** 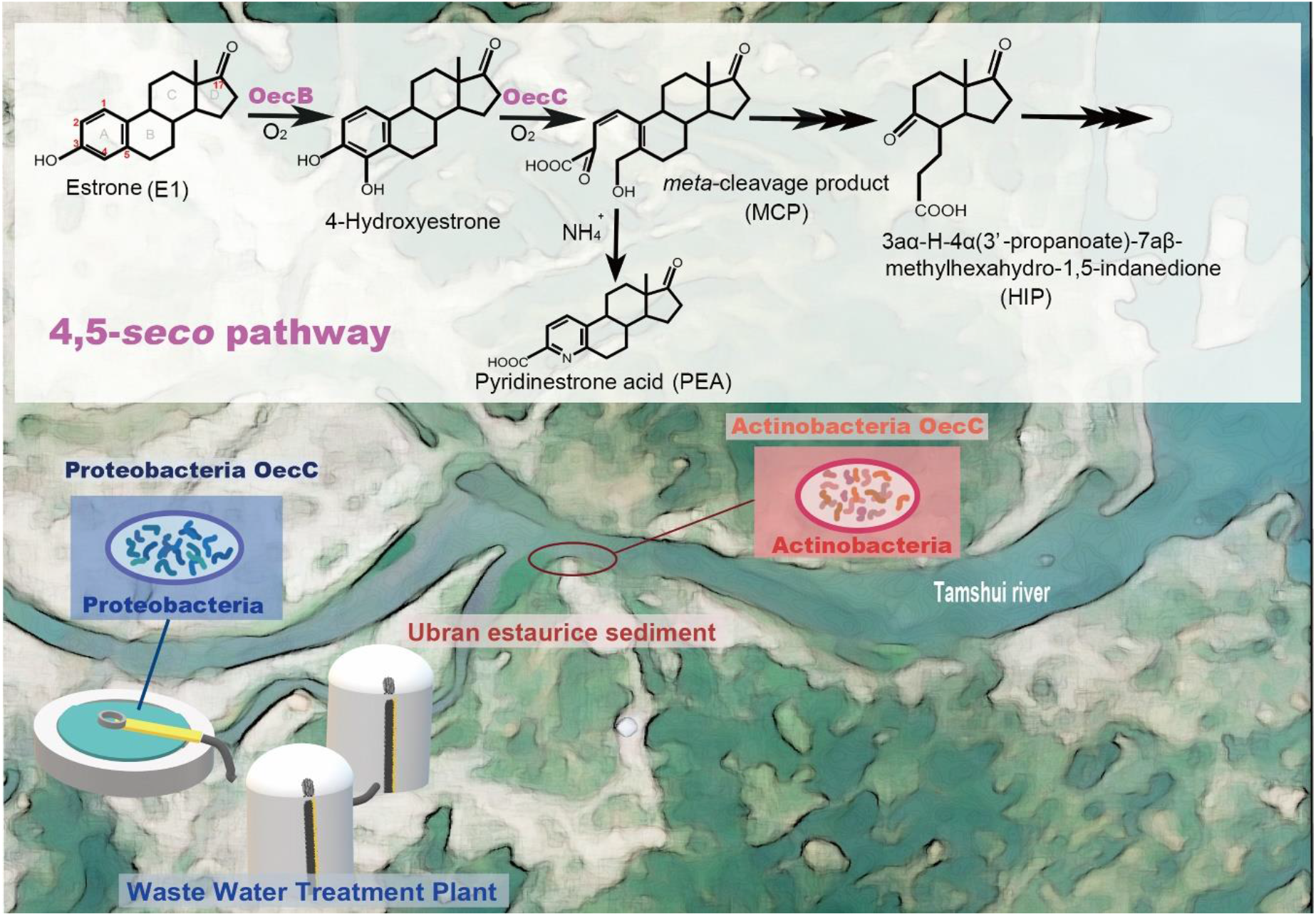

**Highlights:**

- Isolation of an estrogen-degrading actinobacterium *Rhodococcus* sp. strain B50 and establishment of a strain B50 genetic manipulation system.
- Strain B50 exhibits a two-fold estrogen degradation rate of that of estrogen-degrading alpha-proteobacteria under the same cultivation conditions.
- Functional characterization of two oxygenase genes, *oecB* and *oecC*, involved in estrogenic A-ring cleavage in actinobacteria.
- Identification of two extracellular estrogenic metabolites, PEA and HIP, in the estrone-fed strain B50 cultures.
- Detection of actinobacterial *oecC* sequences as well as PEA and HIP in the estrone-spiked urban estuarine sediments.

## 1. Introduction

Estrogens are steroid hormones that regulate the development of the reproductive system and secondary sex characteristics of vertebrates. Natural estrogens include estrone (E1), 17β-estradiol (E2), and estriol (E3). The synthesis and secretion of estrogens exclusively occur in animals, especially in vertebrates (Matsumoto et al., 1997; Tarrant et al., 2003). In the animal liver, estrogens undergo structural modifications (*e.g*., glucuronidation), and are converted into more soluble metabolites to be excreted through urine and feces (Harvey and Farrier, 2011). While required by animals, chronic exposure to trace estrogens at sub-nanomolar levels can disrupt the endocrine system and sexual development in animals (Belfroid et al., 1999; Baronti et al., 2000; Huang and Sedlak, 2001; Kolodziej et al., 2003; Lee et al., 2006). For example, an E2 concentration of 54 ng/L caused severe abnormal development among eelpout embryos (Morthorst et al., 2014). Similarly, the EC50 of E2 causing infertility of fathead minnows (*Pimephales promelas*) was 120 ng/L (Kramer et al., 1998). In addition to being endocrine disruptors, estrogens have been classified as Group 1 carcinogens by the World Health Organization (IARC Monographs-Classifications).

Estrogen pollution has become a global concern and challenge due to the increased human population and mounting demand for livestock products. Livestock manure (Hanselman et al., 2003) and municipal sewage-derived fertilizers (Hamid and Eskicioglu, 2012; Lorenzen et al., 2004) have been a major source of environmental estrogens. Natural estrogens have been considered the most significant contributor to the endocrine-disrupting activity of the swine manure (Noguera-Oviedo and Aga, 2016). However, anaerobic digestion did not alter total estrogen concentrations in livestock manure (Noguera-Oviedo and Aga, 2016), and the estrogens could be released to aquatic ecosystems *via* rainfall and leaching (Hanselman et al., 2003; Kolodziej et al., 2004). Although estrogens could be photodegraded in surface water ecosystems with a degradation half-live ranging from days to weeks (Jurgens et al., 2002; Lin and Reinhard, 2005), photodegradation is hardly occurred in the light-limited environments such as aquatic sediments. As a result, estrogens are often accumulated in urban estuarine sediments downstream to industrialized areas due to their low solubility in water (e.g., 1.5 mg per liter for estradiol) (Shareef et al., 2006) and chemical recalcitrance (Griffith et al., 2016; Wise et al., 2011).

Mineralization of natural estrogens is only accomplished by microorganisms (Thayanukul et al., 2010; Chen et al., 2017 and 2018; Wang et al., 2020; Chiang et al., 2020). Complete estrogen mineralization by bacteria was first described by Coombe *et al*. (1966) in actinobacterium *Nocardia* sp. strain E110. Additionally, *Rhodococcus* isolates (*e.g*., *R. equi* and *R. zopfii*) (Yoshimoto et al., 2004; Kurisu et al., 2010), *Novosphingobium tardaugens* NBRC 16725 (Fujii et al., 2002), and *Sphingomonas* spp. (Ke et al., 2007; Yu et al., 2007) were also capable of mineralizing estrogens. According to current literature, several putative estrogen biodegradation pathways have been proposed (Yu et al., 2013), suggesting that different bacterial taxa likely adopt different degradation strategies to degrade estrogens. Recently, the aerobic 4,5-*seco* pathway for estrogen degradation and the corresponding enzymes in proteobacteria have been studied in some detail (Chen et al., 2017; Wu et al., 2019). However, since common commercial genome-editing techniques are not applicable in these estrogen-degrading alpha-proteobacteria, the function of these enzyme genes remains to be validated. Moreover, homologous genes in the 4,5-*seco* pathway are not found in the genomes of the estrogen-degrading actinobacteria based on sequence homology.

In this study, we used actinobacterium *Rhodococcus* sp. strain B50 isolated from the soil as the model microorganism to study actinobacterial estrogen degradation due to its outstanding efficiency in estrogen degradation and its compatibility with common genetic manipulation techniques: (i) forming independent colonies on agar-based solid media; (ii) incorporating commercial vectors *via* electroporation, and (iii) sensitivity to commercial antibiotics (*e.g*., chloramphenicol). We applied an integrated approach including genomics, metabolomics, and gene disruption experiments to elucidate the estrogen degradation pathway in actinobacteria. Subsequently, we used the extracellular metabolites and the degradation gene *oecC* as biomarkers to investigate estrogen biodegradation in urban estuarine sediment.

## 2. Materials and Methods

### 2.1. Isolation and characterization of *Rhodococcus* sp. strain B50

#### 2.1.1 Enrichment and isolation of strain B50

Soil samples were collected from Dr. Hayashi’s garden in Kodaira, Tokyo, Japan in 2004. To enrich the estrogen-degrading actinobacteria, the soil samples (2 g) were incubated in a rich growth medium (100 mL in a 0.5-L flask) containing (NH_4_)_2_HPO_4_ (0.12 g), KCl (0.25 g), Bacto yeast extract (0.02 g), E1 (0.2 g), and soil extract (20 mL). Medium pH was adjusted to 7.1 with HCl before autoclaving. To prepare the soil extract, soil (500 g) was suspended in double-distilled water (ddH_2_O) (2.4 L) and the soil suspension was autoclaved. After that, the autoclaved soil suspension was centrifuged at 1,000 × *g* for 10 min and the resulting supernatant was defined as the soil extract. The bacterial cultures were incubated at 28°C with continuous shaking (150 rpm) in the dark (to avoid the growth of phototrophs) for 14 days. The E1-spiked enrichment cultures were diluted (10^−4^-fold) and spread on E1-coated agar plates containing (NH_4_)_2_HPO_4_ (0.12 g), KCl (0.25 g), Bacto yeast extract (0.02 g), and soil extract (20 mL). E1 was dissolved in methanol (10 mg/L) and was spread onto the surface of each agar plate; the plates were placed in laminar flow for 2 days at room temperature to remove methanol before inoculation. The plates were then incubated at 28°C for an additional 10 days. Bacterial colonies with a clear zone (in which E1 was exhausted) were selected and streaked on agar plates to obtain single colonies. After a three-day incubation at 28°C, single colonies with a clear zone were further selected and incubated with E1 (1 mM) as the sole carbon and electron donor in a chemically defined mineral medium. The basal medium used for the isolation and routine cultivation of strain B50 contained NH_4_Cl (2.0 g/L), KH_2_PO_4_ (0.67 g/L), and K_2_HPO_4_ (3.95 g/L). After autoclaving, this basal medium was supplemented with MgSO_4_ (2 mM), CaCl_2_ (0.7 mM), filtered vitamin mixture (1,000 x; the stock solution contained cyanocobalamin (50 mg), pantothenic acid (50 mg), riboflavin (50 mg), pyridoxamine (10 mg), biotin (20 mg), folic acid (20 mg), nicotinic acid (25 mg), nicotine amide (25 mg), α-lipoic acid (50 mg), *p*-aminobenzoic acid (50 mg), and thiamine (50 mg) per liter), ethylenediaminetetraacetic acid (EDTA)-chelated trace elements (1,000 x) (Rabus and Widdel, 1995), and sodium selenite (4 μg/L). Dimethyl sulfoxide (DMSO)-dissolved E1 (stock concentration = 125 mM) was added to the mineral medium to a final concentration of 1 mM E1. The 16S rRNA gene sequence was amplified from the total genomic DNA extracted from the bacterial culture using universal primers 27F and 1492R, and the taxonomy of strain B50 was determined using the Nucleotide Basic Local Alignment Search Tool (BLASTn) from the National Center for Biotechnology Information (NCBI).

#### 2.1.2. Aerobic incubation of strain B50 with sex steroids

The wild-type and the gene-disrupted mutants of strain B50 were used in the resting cell biotransformation assays. Bacteria were first aerobically grown in Luria-Bertani (LB) broth (40 mL in a 200-mL Erlenmeyer flask) containing E1 (50 μM) as an inducer at 28°C with continuous shaking (150 rpm). Cells were collected through centrifugation (8,000 × *g*, 20 min, 15°C) at the exponential growth phase with an optical density at 600 nm (OD_600_) of 0.5 (optical path, 1 cm), followed by removal of the supernatant. The cell pellet was resuspended in a chemically defined mineral medium as mentioned above. The cell suspension (OD_600_ = 1; 10 mL) was incubated with E1 or other sex steroids (100 mg/L) and was aerobically incubated at 28°C with continuous shaking (150 rpm) and were sampled (1mL) hourly. The resulting samples were extracted twice using equal volumes of ethyl acetate. The ethyl acetate fractions were evaporated, and the pellets containing estrogen metabolites were stored at -20°C until further analysis.

#### 2.1.3. Extracellular production of PEA and HIP

The preparation of the resting cell assays of strain B50 was the same as described above. The cell suspensions (OD_600_ = 1; 10 mL) were fed with E1 (2-fold serial dilution; from 40 to 1.25 mg/mL) and then aerobically incubated at 28°C with shaking (150 rpm). The cell suspensions were sampled (1 mL) after 24 hours of aerobic incubation with E1. Cells were separated from the growth medium through centrifugation (10,000 × *g*, 10 min) followed by isolation of the supernatant, and the cell pellet was discarded. To facilitate the extraction of PEA and HIP, the samples were acidified using 30 μL of 6N HCl. The resulting samples were extracted twice using ethyl acetate (1 mL), and the extracted estrogen metabolites were stored at -20°C until further analysis.

### 2.2. *oecB* and *oecC* disruption in strain B50

The disruption of *oecB* and *oecC* in strain B50 were performed using homologous recombination by a pK18-Cm^R^-pheS** plasmid (**Fig. 3A**). First, gene-specific DNA fragments, including the 900 base pair flanking region and the 100 base pair coding sequence, of the target genes were cloned through Platinum polymerase (Thermo Fisher Scientific) and purified with GenepHlow^™^ Gel/PCR kit (Geneaid). The up-stream, down-stream, and plasmid backbone fragments were assembled *via* an In-Fusion® HD Cloning Kit (TAKARA Bio; Kusatsu, Shiga, Japan) to generate the plasmid (*oecB*- or *oecC*-pK18-Cm^R^-pheS**). This plasmid was electroporated into *E. coli* strain S17-1 using a Gene Pulser Xcell^™^ (Bio-Rad, CA, USA) with the conditions of 2.5kV, 25μF, and 200Ω. The transformed *E. coli* strain S17-1 was co-incubated with wild-type *Rhodococcus* sp. strain B50 at 30°C overnight for horizontal gene transfer through conjugation. Successfully transformed colonies of *Rhodococcus* sp. strain B50 were selected with nalidixic acid and chloramphenicol. The insertion of the chloramphenicol-resistant gene into the strain B50 genome was confirmed using the plasmid-specific primer pairs 5′-TTCATCATGCCGTTTGTGAT-3′ (Pk18-cmpheS-F) and 5′-ATCGTCAGACCCTTGTCCAC-3′ (Pk18-cmpheS-R). The genotypes of the *oecB*- and *oecC*-disrupted mutants were further examined using the gene-specific primer pairs [5′-AGGTCGATGTCCTCGACACCGAGG-3′ (*oecB*-3k-F) and 5′-CGCATCCTCAGTCACCTCGGCG-3′ (*oecB*-3k-R)] and [′’-AACCATGATCTTCACCATCG-3′ (*oecC*-3k-F) and 5′-TCAGTAGCCGTGCACGAG-3′ (*oecC*-3k-R)], respectively.

**Figure 1.**
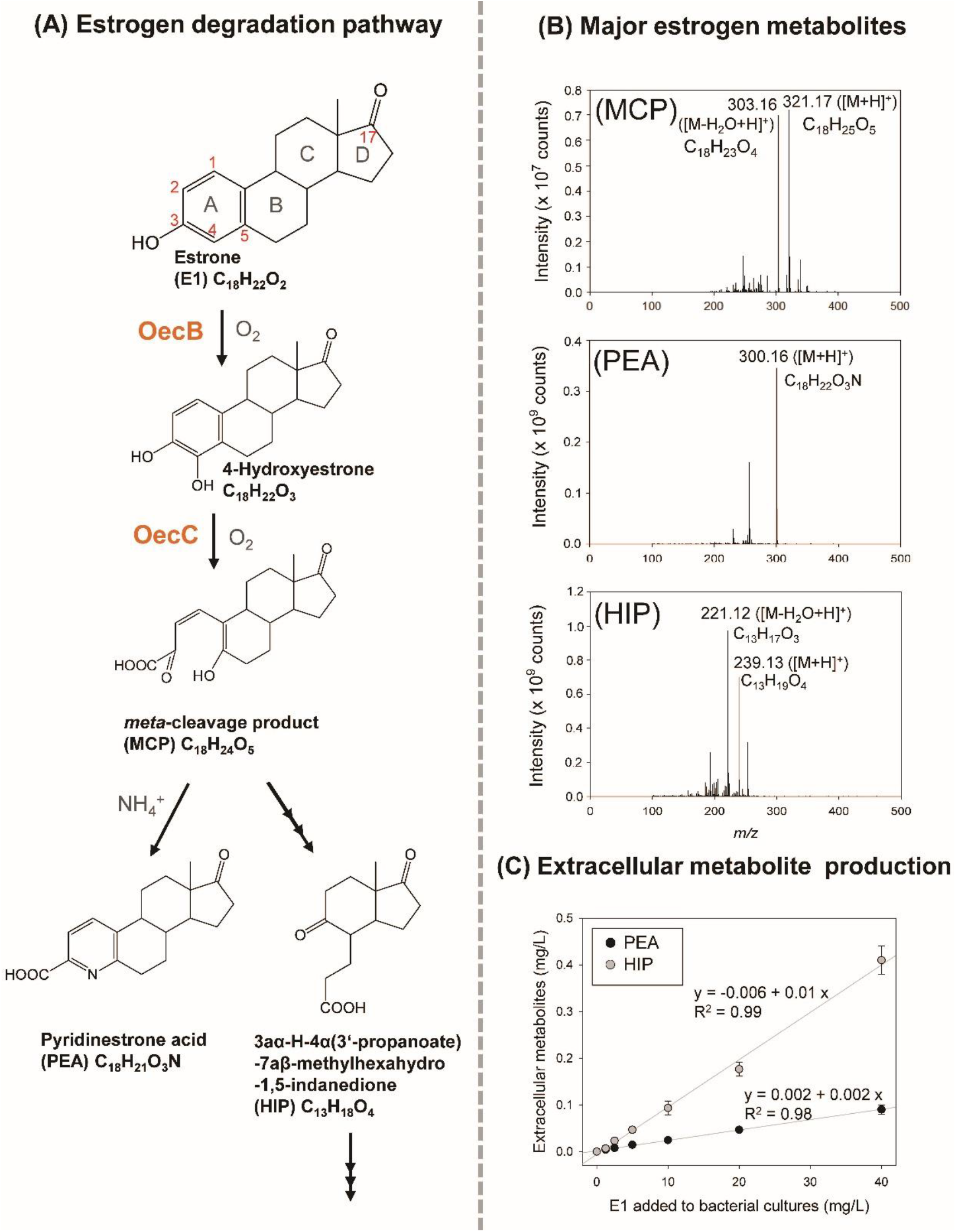
Metabolites and genes involved in estrogen degradation of *Rhodococcus* sp. strain B50. (**A**) The proposed pathway for estrogen degradation of strain B50. OecB, estrone 4-hydroxylase; OecC, 4-hydroxyestrone 4,5-dioxygenase. (**B**) Mass spectrometry spectra of major estrogen metabolites of strain B50 incubated with E1 (100 mg/L). The UPLC and MS behaviors of these metabolites are shown in **Table S2**. (**C**) The dose-dependent manner of the extracellular metabolite production in the E1-grown strain B50 cultures.

**Figure 2.**
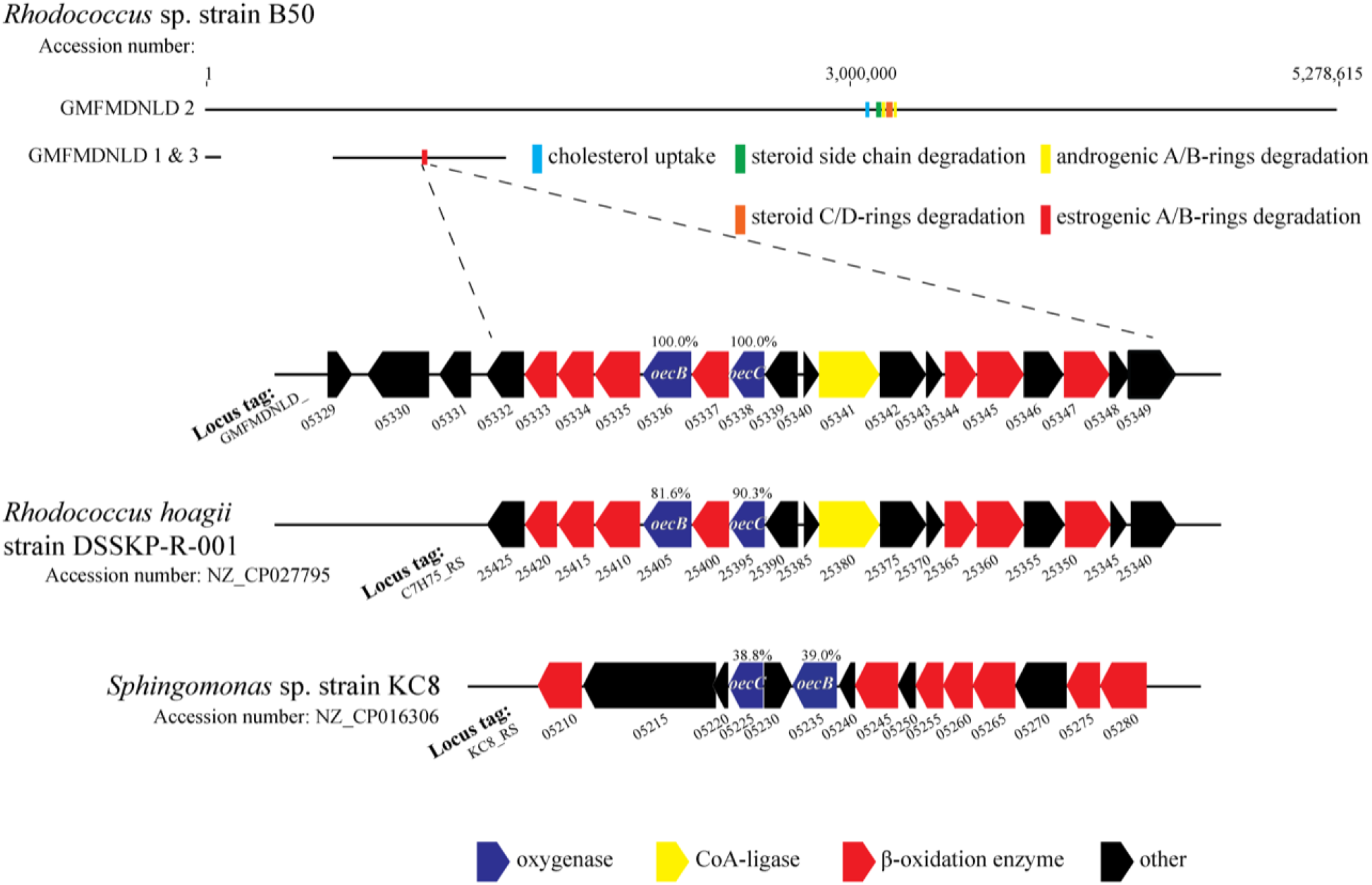
Comparative genomic analysis of *Rhodococcus* sp. strain B50. Gene clusters for cholesterol uptake (blue), cholesterol side-chain degradation (green), androgenic A/B-rings degradation (yellow), and HIP degradation (orange) are located in the linear chromosome, whereas the gene cluster specific for estrogenic A/B-rings degradation (red) is present in the megaplasmid. Percentage (%) indicates the shared identity of the deduced amino acid sequences of the two oxygenase genes.

**Figure 3.**
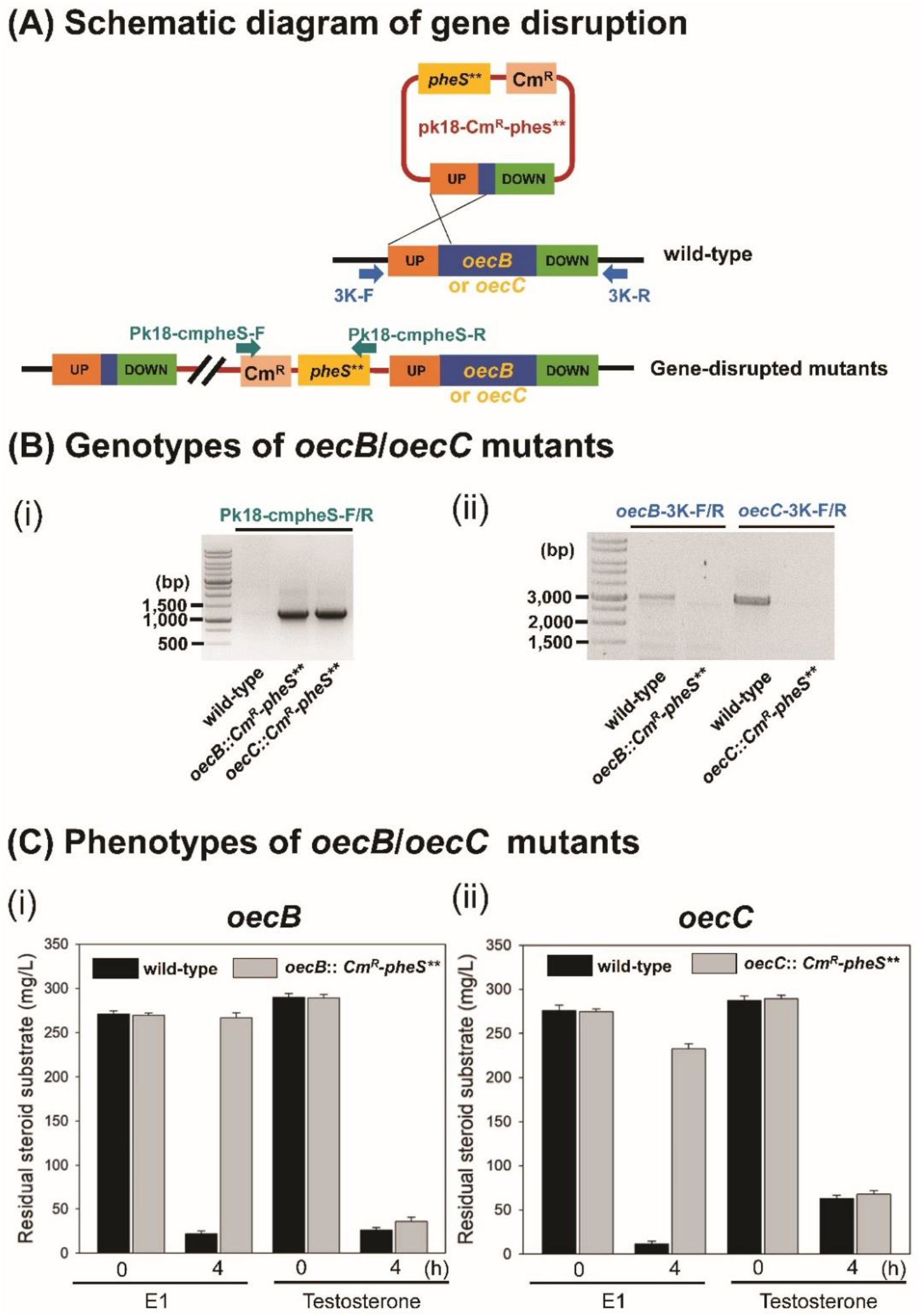
Disruption of *oecB* and *oecC* in strain B50. (**A**) Schematic diagram of homologous recombination-mediated gene disruption. (**B**) Genotype examinations of *oecB*- and *oecC*-disrupted strain B50 mutants. (**Bi**) Agarose gel electrophoresis indicated the insertion of a chloramphenicol-resistant gene (Cm^R^) and *pheS*\*\* cassette into the target genes. (**Bii**) Agarose gel electrophoresis confirmed the gene disruption of *oecB* and *oecC*. (**C**) Phenotypes of *oecB*- and *oecC*-disrupted strain B50 mutants. The wild-type strain B50 was also tested for a comparison. Data shown are the means ± S.D. of three experimental replicates.

### 2.3. Environmental estrogen biodegradation

#### 2.3.1. Sampling site and estuarine sediment sample collection

Approximately six million people reside in the basin of the Tamsui River, Taipei, Taiwan. The Tamsui River estuary receives sewage discharge and waste effluent from the Taipei metropolitan area (Kao et al., 2013). This includes effluent from the Dihua Sewage Treatment Plant, which contains estrogens (approximately 1 ng/L) (Chen et al., 2018). Our sampling site, Guandu (25°6′59.56″N, 121°27′46.99″E), with a salinity of 5–22 parts per thousand (ppt) (Kao et al., 2013; Shih et al., 2017), is located in the upper estuary where the Keelung River meets the main channel of the Tamsui River, and the sewage discharge and seawater intrusion are mixed in Guandu (Kao et al., 2013). In the current study, four sediment cores were collected from Guandu. The sediment samples were collected using polyvinyl chloride corers (7.5-cm diameter). During low tide on May 20, 2019, the corers were pressed down approximately 30 cm into the sediments and sealed with a rubber stopper immediately after collection. Estuarine water samples (20 L) were collected from Guandu on the same day. The sediment and estuarine water samples were carried to the laboratory within 1 hour after sampling and were processed immediately.

#### 2.3.2. Spiking the estuarine sediments with E1

Each sediment core was fractionated into three sections: a subsurface layer (0–5 cm depth), a middle layer (6–10 cm depth), and a bottom layer (11–15 cm depth). Vertical sectioning of the sediment cores was based on the vertical distributions of chemicals and bacteria in the Guandu sediments (Shih et al., 2017). The subsurface layer sediment (1 g) was added to 100-mL Erlenmeyer flasks containing river water (9 mL). The microcosms (10 mL) were then spiked with [3,4C-^13^C]E1 (10 μg/mL) and incubated in the dark at 30°C with stirring (150 rpm). Estrogen metabolites in the microcosms were sampled every two days (0–10 days) and were detected using ultra-performance liquid chromatography–atmospheric pressure chemical ionization–high resolution mass spectrometry (UPLC−APCI−HRMS). The microcosms were also sampled (1 mL) every hour (0–8 hours) and stored at −80°C before the RNA extraction. Estrogen metabolites in the samples were detected using UPLC−APCI−HRMS. The functional *oecC* genes in the microcosm samples were analyzed through PCR-based functional assays as described below.

#### 2.3.3. RNA isolation and cDNA preparation

Total RNA was extracted from the E1-spiked estuarine sediment sample using the RNeasy® PowerSoil® total RNA kit (Qiagen, Hilden, Germany). The crude total RNA was further purified using Turbo DNA-free Kit (Thermo Fisher Scientific) to remove DNA. The DNA-free total RNA was reverse-transcribed to cDNA using the SuperScript^®^ IV First-Strand Synthesis System (Thermo Fisher Scientific) with random hexamer primers (Thermo Fisher Scientific).

#### 2.3.4. Amplification of *oecC* genes from the estuarine sediment samples using degenerate primers

Multiple alignments of *oecC* genes from estrogen-degrading actinobacteria or alpha-proteobacteria were conducted with Geneious® 11.1.5 (Biomatters; Auckland, New Zealand). Degenerate primer pairs were designed according to the conserved regions of actinobacteria (forward: 5′-CGYGGCATCGGATACATCGG-3′; reverse: 5′-ACMGGGTCGCAKCCGATCTC-3′) or alpha-proteobacteria (forward: 5′-CDGYYTGGGCTATSTSGG-3′; reverse: 5′-ATCGCGYCSCASCCRATYTC-3′) respectively. The *oecC* fragments were amplified with PCR with a program of 95°C for 1 min, followed by 30 cycles at 95°C for 30 s, 64°C for 30 s, 72°C for 60 s, and finally 72°C for 5 min. Amplified *oecC* sequences were cloned into *E. coli* DH5α-derived ECOS^™^ 101 competent cells (Yeastern Biotech; Taipei, Taiwan) using the yT&A Cloning Kit (Yeastern Biotech; Taipei, Taiwan). The *oecC* fragments (approximately 800 bp) were sequenced on an ABI 3730xI DNA Analyzer (Applied Biosystems; Waltham, MA, USA) with the BigDye Terminator kit according to the manufacturer’s instructions by the DNA Sequencing Core Facility at Academia Sinica.

Other materials and methods for general chemical analyses and molecular biological manipulation are described in **Supplemental Materials and Methods**.

## 3. Results

### 3.1. Isolation and characterization of the estrogen-degrading *Rhodococcus* sp. strain B50

Strain B50 was isolated from a garden soil sample in Tokyo, Japan. The E1-degrading actinobacterium was highly enriched by repeating 10^−8^ dilution transfers. The highly enriched culture was then spread on an agar-based solid medium. Subsequently, independent colonies were selected for incubation in a chemically defined mineral medium containing E1 (270 mg/L) as the sole carbon source and electron donor to confirm their capability to degrade estrogens. We isolated four estrogen-degrading *Rhodococcus* spp. strains, including strain B50, from the soil samples (see **Fig. S1** for the morphology of bacterial cells and colonies of strain B50). Among them, strain B50 is inherently sensitive to various antibiotics including ampicillin, kanamycin, and chloramphenicol (**Table S1**), as opposed to the other three strains. Thus, strain B50 was selected as the model microorganism to study actinobacterial estrogen biodegradation for its convenience in antibiotics-based genetic manipulation. All of the *Rhodococcus* spp. isolates are resistant to the quinolone antibiotic nalidixic acid, which enables plasmid transfer from *E. coli* (nalidixic acid-sensitive) through conjugation and allows the selective growth of *Rhodococcus* strains. Next, we characterized the substrate spectra of strain B50. Strain B50 can utilize E1, E2, E3, testosterone, or cholesterol as the sole carbon source and electron donor (**Fig. S2**). The doubling time of strain B50 which grows on E1, testosterone, and cholesterol ranges from 3–4, 6–8, and 12–14 hours, respectively.

### 3.2. Identification of estrogen metabolites

The growth experiments suggested that strain B50 can mineralize estrogens. To characterize the estrogen degradation pathway in strain B50, strain 50 resting cells (∼10^9^ cells/mL) were aerobically incubated with E1 (10 mg/L), sampled hourly, extracted using ethyl acetate, and the metabolite profile was analyzed through UPLC**–**APCI**–**HRMS. The metabolite profile analysis revealed at least four E1-derived metabolites, including PEA and HIP in the established 4,5-*seco* pathway (**Table S2**). The retention time of the detected metabolites in the UPLC and their HRMS behaviors were identical to those of the corresponding authentic standards (**Fig. 1B** and **Table S2**), suggesting that strain B50 adopts the 4,5-*seco* pathway to degrade estrogens. Moreover, we observed the accumulation of both PEA and HIP in the supernatants of strain B50 cultures in a dose-dependent manner depending on added E1 (**Fig. 1C**).

### 3.3. Identification of the estrogen-degrading genes *via* comparative genomic analysis

Metabolite profile analysis suggested that strain B50 degrades estrogens *via* the 4,5-*seco* pathway established in proteobacteria. However, the homologous genes involved in the proteobacterial 4,5-*seco* pathway were not annotated in the strain B50 genome, likely due to distant phylogeny between proteobacteria and actinobacteria. Therefore, we compared the strain B50 genome to the genomes of the reported estrogen-degrading actinobacteria in the database. Through the comparative genomic analysis, we identified a putative estrogen-degrading gene cluster (GMFMDNLD _05329 to 05349; **Dataset S1**) on a circular genetic element (i.e., megaplasmid; GMFMDNLD 3) of strain B50, which is also present in the genome of estrogen-degrading *Rhodococcus* sp. strain DSSKP-R-001 (Zhao et al., 2018), but not in other *Rhodococcus* members incapable of degrading estrogen. Moreover, the two homologous estrogen-degrading gene clusters are both located on their megaplasmids (**Fig. 2; Dataset S1**). Among them, the gene cluster of strain B50 is surrounded by a transcriptional regulator and a transposase gene (GMFMDNLD _05329 and 05330). In the putative estrogen-degrading gene cluster, GMFMDNLD _05338 encodes a putative *meta*-cleavage enzyme, which likely functions as the 4-hydroxyestrone 4,5-dioxygenase (OecC; Chen et al., 2017). Moreover, GMFMDNLD_05336 encodes a member of the cytochrome P450 protein family and thus likely functions as an oxygen-dependent estrone 4-hydroxylase (OecB). The nucleotide sequences of 16S rRNA, and the *oecB* and *oecC* genes of strain B50 are shown in **Appendix S1, S2, and S3**, respectively.

### 3.4. Functional validation of actinobacterial *oecB* and *oecC* in estrogenic A-ring degradation

Next, we aimed to confirm the function of the putative oxygenase genes *oecB* and *oecC* involved in actinobacterial estrogen degradation. Thus, we disrupted the putative *oecB* (GMFMDNLD_05336) and *oecC* (GMFMDNLD_05338) in strain B50 using site-directed mutagenesis (insertion of a chloramphenicol-resistance gene (Cm^R^) and *pheS*\*\* cassette). The plasmid was transferred from *E. coli* (nalidixic acid-sensitive) to strain B50 (nalidixic acid-resistant) through conjugation. Then, the *oecB*- and *oecC*-disrupted strain B50 mutants were selected and maintained on LB agar containing two antibiotics: chloramphenicol (25 μg/mL) and nalidixic acid (12.5 μg/mL). PCR with primers flanking the *oecB* and the Cm^R^ genes confirmed successful insertion of the chloramphenicol-resistance cassette into the *oecB* gene in the mutated strain (**Fig. 3B**). The *oecB*-disrupted mutant can utilize testosterone but not E1 (**Fig. 3Ci**), while the wild-type strain B50 apparently degraded testosterone and E1 within 4 hours. Moreover, we did not observe any estrogenic metabolites [*e.g*., 4-hydroxyestrone, the *meta*-cleavage product (MCP), PEA, or HIP] in the *oecB*-disrupted strain B50 mutant incubated with E1, suggesting that *oecB* is involved in the transformation of E1 into 4-hydroxyestrone (**Fig. S3**). Applying the same gene-disruption approach (insertion of the Cm^R^ and *pheS*\*\* cassette), we obtained an *oecC*-disrupted strain B50 mutant (**Fig. 3B**). Similarly, the *oecC*-disrupted mutant can only utilize testosterone but failed to utilize E1 (**Fig. 3Cii**). Moreover, we observed 4-hydroxyestrone accumulation but no downstream products in the *oecC*-disrupted strain B50 mutant cultures incubated with E1 (**Fig. S3**), revealing that *oecC* is involved in estrogenic A-ring cleavage.

### 3.5. Alignment of the *oecC* genes from actinobacteria and proteobacteria

Our data suggest that both proteobacteria and actinobacteria adopt the *meta*-cleavage dioxygenase OecC to cleave the A-ring of natural estrogens. The phylogenetic tree shows that *oecC* orthologs from all known estrogen-degrading bacteria in the database form a distinct lineage (**Fig. S4**), separated from the *hsaC* and *tesB*, which are involved in androgenic A-ring cleavage in bacteria (**Fig. S5**). Proteobacteria-specific *oecC* primers have been designed and examined in our previous study (Chen et al., 2018). In the present study, we aimed to design specific primers for actinobacterial *oecC*. The phylogenetic divergence of *oecC* sequences between actinobacteria and proteobacteria allows the design of taxa-specific primers for environmental studies (**Fig. 4A**). The designed actinobacterial primers were validated using chromosomal DNA of the three other estrogen-degrading *Rhodococcus* spp. strains isolated as mentioned in **3.1**. To test primer specificity, gDNA from an estrogen-degrading proteobacterium *Sphingomonas* sp. strain KC8 and from a testosterone-degrading actinobacterium *Gordonia cholesterolivorans* incapable of degrading estrogens were used as negative controls. PCR products with an expected size of approximately 800 base pairs were only amplified from gDNA of the estrogen-degrading *Rhodococcus* spp. but not from gDNA of *G. cholesterolivorans* or strain KC8 (**Fig. 4B**), suggesting that the degenerate *oecC* primer is highly specific to actinobacterial *oecC* and cannot be used to amplify the androgenic *meta*-cleavage dioxygenase gene *hsaC* and proteobacterial *oecC*.

**Figure 4.**
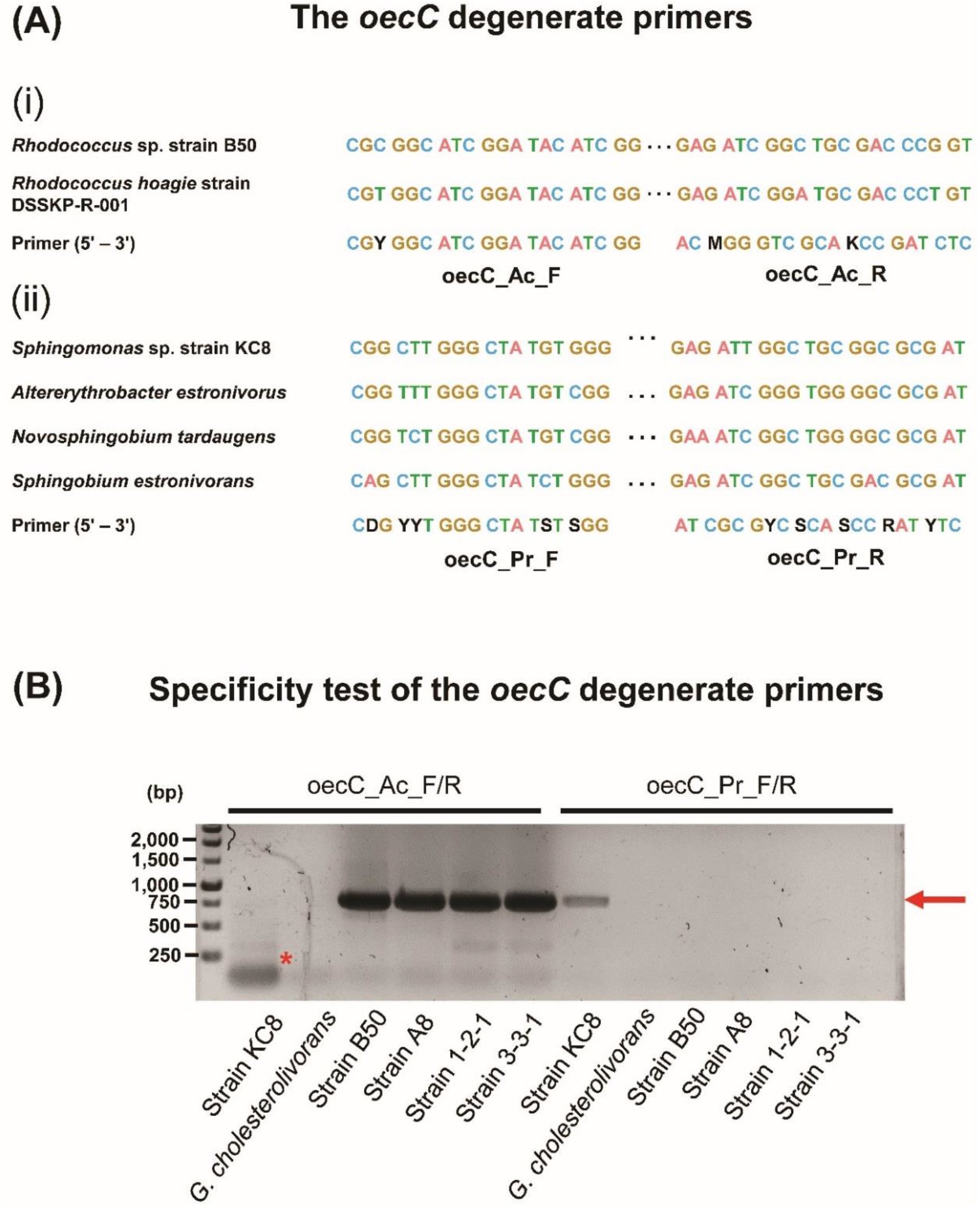
PCR-based functional assay using degenerate primers derived from *oecC* genes. (**A**) Highly conserved nucleotide sequence regions of *oecC* derived from (**Ai**) actinobacteria and (**Aii**) proteobacteria used to deduce degenerate primers. (**B**) Agarose gel electrophoresis indicated that *oecC* is harbored only by estrogen-degrading *Rhodococcus* spp. but not by testosterone-degrading actinobacterium *Gordonia cholesterolivorans* strain DSM 45229 or estrogen-degrading alpha-proteobacterium *Sphingomonas* sp. strain KC8. R = A + G, S = G + C, and Y = C + T. Arrow indicates the expected fragment of *oecC*; asterisk demonstrates the primer dimer.

### 3.6. The metabolite profile and *oecC*-based functional analyses reveals actinobacteria as active estrogen degraders in urban estuarine sediment

Subsequently, the actinobacterial and proteobacterial *oecC* degenerate primers were used to study estrogen biodegradation in the urban estuarine sediment of the Tamsui River, a river passing through the Taipei metropolitan area in Taiwan. [3,4C-^13^C]E1 (100 μg/g sediment) was spiked into the urban estuarine sediment samples. Metabolite profile analysis revealed time-dependent PEA and HIP accumulation in the supernatants of the sediment samples, suggesting the occurrence of estrogen degradation in the sediment samples (**Fig. 5**). Moreover, a higher concentration of HIP (2 μg/g sediment) was produced by sediment microbiota after 8 days of incubation with [3,4C-^13^C]E1, compared to that of PEA (0.2 μg/g sediment).

**Figure 5.**
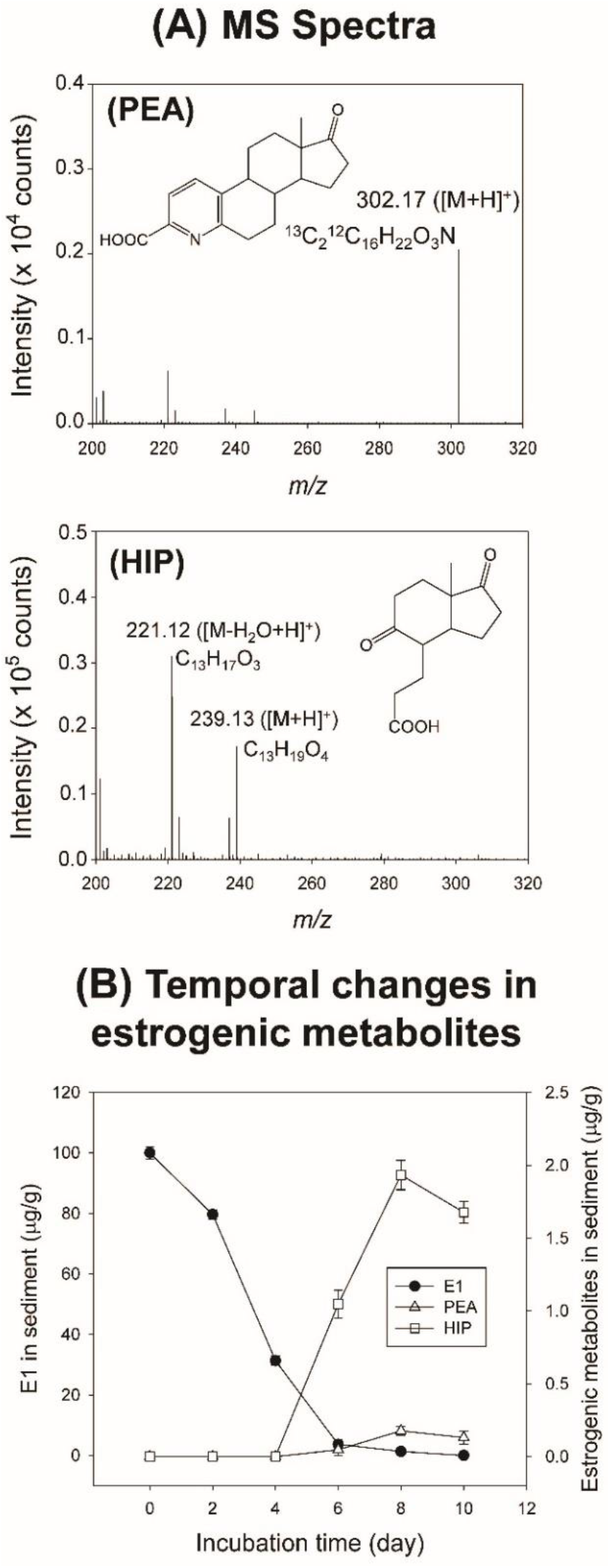
UPLC–APCI–HRMS detection of PEA and HIP in [3,4C-^13^C]E1 (100 μg/g sediment)-spiked estuarine sediment. (**A**) MS spectra of the estrogenic metabolites PEA and HIP. (**B**) Temporal change in estrogenic metabolite production. Measurements of PEA and HIP were based on the adducts corresponding to the two compounds using UPLC–APCI–HRMS. Data shown are the means ± S.D. of three experimental replicates.

Total RNA was extracted and purified from the [3,4C-^13^C]E1-spiked sediment samples hourly. Reverse transcribed cDNA was used as the template for the *oecC* degenerate primers in the PCR-based assays. After an eight-hour incubation with [3,4C-^13^C]E1, we detected the *oecC* amplicons in the PCR experiment using the actinobacterial *oecC* primers but not in the experiment using the proteobacterial *oecC* primers (**Fig. 6A**). Next, the actinobacterial *oecC* amplicons were cloned into *E. coli* strain DH5α. Ten clones (sediment cDNA #1–10) were randomly selected for sequencing (**Appendix S4**). Notably, all of the ten *oecC* amplicon sequences obtained from the [3,4C-^13^C]E1-spiked sediment samples were highly similar to that of strain B50 *oecC* (**Fig. 6B**) but were distant from the proteobacterial *oecC* sequences. Altogether, our functional assays suggest that actinobacteria, rather than proteobacteria functioning in the wastewater treatment plants, are responsible for estrogen biodegradation in urban estuarine sediment.

**Figure 6.**
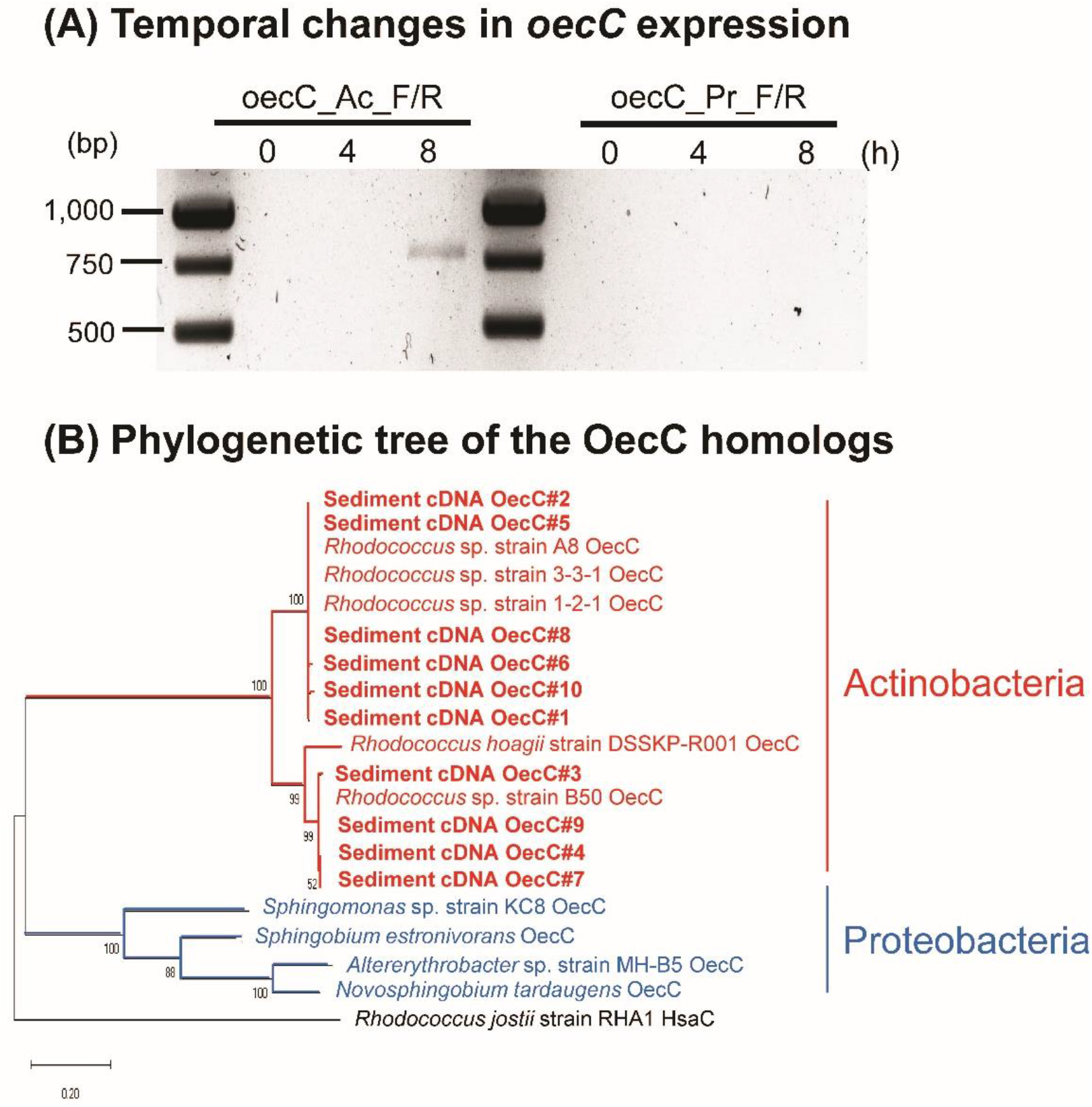
Phylogenetic identification of *oecC* expressed in the [3,4C-^13^C]E1-spiked estuarine sediments. (**A**) RT-PCR indicated temporal changes in the expression of the actinobacterial and proteobacterial *oecC* genes in [3,4C-^13^C]E1-spiked estuarine sediment. (**B**) Phylogenetic tree of deduced amino acid sequences of the *oecC* genes from estrogen-degrading bacterial isolates and *oecC* fragments obtained from the cDNA of the [3,4C-^13^C]E1-spiked estuarine sediment.

## 4. Discussion

### 4.1. A shared estrogen degradation pathway in both actinobacteria and proteobacteria

In the present study, the identification of the estrogenic metabolites PEA and HIP, along with the identification of degradation genes *oecB* and *oecC*, in strain B50 reveals that actinobacteria also adopt the 4,5-*seco* pathway to degrade natural estrogens. Actinobacteria such as *Mycobacterium* spp. and *Rhodococcus* spp. use the flavin-dependent monooxygenase *hsaAB*, with *hsaA* and *hsaB* as the oxygenase and reductase subunit, respectively, to add a hydroxyl group to the C-4 of the androgenic metabolite 3-hydroxy-9,10-seconandrost-1,3,5(10)-triene-9,17-dione (3-HSA) (**Fig. S5**) (Dresen et al., 2010; Bergstrand et al., 2016; Holert et al., 2018). Therefore, in a previous study using the *Sphingomonas* sp. strain KC8 as the model organism (Chen et al., 2017), we speculated that a similar *hsaA*-type gene might be responsible for the transformation of E1 into 4-hydroxyestrone in strain KC8. In the present study, we demonstrated that the functionally confirmed *oecB* of strain B50 is a heme-dependent cytochrome P450-type monooxygenase gene but not a flavin-dependent *hsaA*-type monooxygenase gene. Surprisingly, a highly similar gene is also present in the *Sphingomonas* sp. strain KC8 as well as other estrogen-degrading proteobacteria. Thus, our data suggest that both proteobacteria and actinobacteria use the cytochrome P450-type monooxygenase, but not the putative flavin-dependent monooxygenase, to transform E1 into 4-hydroxyestrone. The degradation rate of strain B50 (approximately 4.5 mg E1/h) is almost two times faster than that of alpha-proteobacterium *Sphingomonas* sp. strain KC8 (approximately 2.5 mg E1/h) (Chen et al., 2017) under similar cultivation conditions, suggesting that actinobacteria are more efficient in estrogen degradation.

### 4.2. Identification of the specific gene cluster for aerobic degradation of estrogenic A/B-rings

In the strain B50 genome, we identified a gene cluster responsible for estrogen degradation. The disruption of the oxygenase genes *oecB* and *oecC* in this gene cluster apparently aborted estrogen degradation by strain B50, indicating direct involvement of *oecB* and *oecC* in the activation and cleavage of the estrogenic A-ring. The *oecB* is a typical cytochrome P450 monooxygenase which requires O_2_ and NADPH as the co-substrate and reductant, respectively. Only a minor amount of the expected metabolite 4-hydroxyestrone was produced from E1 in the bacterial cultures of the *oecC* mutant, likely related to the shortage of NADPH regeneration in strain B50 cells as E1 cannot be oxidized by the *oecC*-disrupted mutant to generate reducing equivalents. However, the function of the other genes in this gene cluster remains to be investigated through gene-disruption experiments and phenotype analysis. For example, after the estrogenic A-ring cleavage, the *meta*-cleavage product should be further degraded by the gene products in this gene cluster, such as the putative CoA-ligase and two sets of β-oxidation enzymes. Moreover, this gene cluster is located in the megaplasmid of strain B50 and is surrounded by transposon elements and transposases. In parallel, a highly similar gene cluster (C7H75_25375∼_25425) is also present in a plasmid (plas2; 95,132 bp) of the estrogen-degrading *Rhodococcus* sp. strain DSSKP-R-001 (Zhao et al., 2018). The coincidence raises speculation that the estrogen-degrading capacity in the environment may be conferred among actinobacteria via horizontal gene transfer. This gene cluster may thus be used as a biomarker to identify actinobacteria capable of estrogen degradation. Therefore, one may assess the estrogen degradation potential of various actinobacterial strains in different environments by probing this gene cluster in the metagenomic and/or metatranscriptomic data.

### 4.3. The extracellular metabolites PEA and HIP are competent biomarkers for assessing the occurrence and fate of estrogen in environmental samples

A highlight in this study is the detection of two extracellular metabolites (i.e., PEA and HIP) in strain B50 cultures with added E1. Bacteria depend on HIP-CoA ligase (FadD3) to activate HIP, enabling further degradation of the estrogenic C/D-rings (Crowe et al., 2018; Wu et al., 2019). CoA is an essential cofactor in numerous biosynthetic and energy-yielding metabolic pathways (Boll et al., 2020). When CoA is required in other metabolic pathways, the CoA-esters in the 4,5-*seco* pathway (e.g., HIP-CoA) can be deconjugated (Takamura and Nomura, 1988; Lin et al., 2015). The deconjugated metabolites like HIP are often toxic to bacterial cells, and are therefore excreted to the medium (Wu et al., 2019). Our data revealed that approximately 0.2∼0.5% and 1∼2 % of E1 molecules are transformed to PEA and HIP during estrogen degradation by strain B50, respectively. We thus propose that PEA and HIP may be suitable biomarkers for monitoring environmental estrogen biodegradation because (**i**) these two metabolites are produced by two major estrogen-degrading bacterial taxa, namely actinobacteria and proteobacteria; (**ii**) PEA and HIP are critical metabolites for estrogenic A-ring and B-ring degradation, respectively; (**iii**) the extracellular accumulation of these two metabolites exhibited an estrogen dose-dependent manner; (**iv**) these two metabolites can be easily detected using UPLC-HRMS (detection limits at picomolar level); and (**v**) PEA is exclusively produced during bacterial estrogen degradation.

## 5. Conclusion

In summary, we identified extracellular metabolites (PEA and HIP) and two essential genes (*oecB* and *oecC*) involved in actinobacterial estrogen degradation. Since the phenolic A-ring of steroidal estrogens directly interacts with the estrogen receptor (Baker and Lathe, 2018), the cleavage of the phenolic A-ring eliminates their estrogenic activity (Chen et al., 2017). Therefore, PEA and HIP, along with *oecC*, are suitable biomarkers for monitoring the water quality of environments contaminated by estrogens. We designed and tested specific primers for *oecC* of proteobacteria (Chen et al., 2018) and actinobacteria (this study). Interestingly, while previous studies suggested that proteobacteria are the major estrogen consumers in wastewater treatment plants, our PCR-based functional assays demonstrate that actinobacteria are active estrogen degraders in urban estuarine sediments. The combination of targeted metabolites analysis with PCR-based functional assays thus represents a simple, cost-effective, and rapid approach to gain a holistic view of the fate of steroidal estrogens in the environment. Nevertheless, the contribution of proteobacteria in this estrogen-contaminated aquatic ecosystem could be underestimated due to the potential bias of the *oecC*-specific primers. Finally, the gene cluster containing the estrogen-degrading geneses in strain B50 and strain DSSKP-R-001 are all present in their plasmids. Therefore, the *oecB*- and *oecC*-containing plasmids could also be used to transform other actinobacteria into efficient estrogen degraders or even with a broader substrate spectrum via gene knock-in.

## Acknowledgments

This study was supported by the Ministry of Science and Technology of Taiwan (109-2221-E-001-002). Po-Hsiang Wang was supported by the Research and Development Office as well as Research Center for Sustainable Environmental Technology, National Central University, Taiwan. Yi-Lung Chen was supported by the Research Grants for New Teachers of College of Science, Soochow University, Taiwan. We thank the Institute of Plant and Microbial Biology, Academia Sinica, for providing access to the Small Molecule Metabolomics Core Facility (for UPLC–HRMS analyses).

## Author Contributions

Y.-R.C. and P.-H.W. designed the research. T.-H. H., Y.-L.C., and M.-R.C. performed the research. M..M., M.H., and T.H. contributed new reagents and analytic tools. Y.-L.C., T.-H.H., and Y.-R.C analyzed the data. Y.-R.C and P.-H.W. drafted the manuscript. All authors reviewed the manuscript.

## References

Baker, M.E., Lathe, R., 2018. The promiscuous estrogen receptor: Evolution of physiological estrogens and response to phytochemicals and endocrine disruptors. J. Steroid Biochem. Mol. Biol. 184, 29–37.

Baronti, C., Curini, R., D’Ascenzo, G., Di Corcia, A., Gentili, A., Samperi, R., 2000. Monitoring Natural and Synthetic Estrogens at Activated Sludge Sewage Treatment Plants and in a Receiving River Water. Environ. Sci. Technol. 34, 5059–5066.

Belfroid, A.C., Van der Horst, A., Vethaak, A.D., Schäfer, A.J., Rijs, G.B., Wegener, J., Cofino, W.P., 1999. Analysis and occurrence of estrogenic hormones and their glucuronides in surface water and waste water in The Netherlands. Sci. Total Environ. 225, 101–108.

Bergstrand, L.H., Cardenas, E., Holert, J., Van Hamme, J.D., Mohn, W.W., 2016. Delineation of steroid-degrading microorganisms through comparative genomic analysis. mBio. 7, e00166.

Boll, M., Geiger, R., Junghare, M., Schink, B., 2020. Microbial degradation of phthalates: biochemistry and environmental implications. Environ. Microbiol. Rep. 12, 3–15.

Chen, Y.-L., Fu, H.-Y., Lee, T.-H., Shih, C.-J., Huang, L., Wang, Y.-S., Ismail, W., Chiang, Y.-R., 2018. Estrogen Degraders and Estrogen Degradation Pathway Identified in an Activated Sludge. Appl. Environ. Microbiol. 84, e00001–18.

Chen, Y.-L., Yu, C.-P., Lee, T.-H., Goh, K.-S., Chu, K.-H., Wang, P.-H., Ismail, W., Shih, C.-J., Chiang, Y.-R., 2017. Biochemical Mechanisms and Catabolic Enzymes Involved in Bacterial Estrogen Degradation Pathways. Cell Chem. Biol. 24, 712–724.

Chiang, Y.-R., Wei, S.T.-S., Wang, P.-H., Wu, P.-H., Yu, C.-P., 2020. Microbial degradation of steroid sex hormones: implications for environmental and ecological studies. Microb. Biotechnol. 13, 926–949.

Coombre, R.G., Tsong, Y.Y., Hamilton, P.B., Sih, C.J., 1966. Mechanisms of steroid oxidation by microorganisms. X. Oxidative cleavage of estrone. J. Biol. Chem. 241, 1587–1595.

Crowe, A.M., Workman, S.D., Watanabe, N., Worrall, L.J., Strynadka, N.C.J., Eltis, L.D., 2018. IpdAB, a virulence factor in Mycobacterium tuberculosis, is a cholesterol ring-cleaving hydrolase. Proc. Natl. Acad. Sci. USA. 115, E3378–E3387.

Dresen, C., Lin, L.Y.-C., D’Angelo, I., Tocheva, E.I., Strynadka, N., Eltis, L.D., 2010. A flavin-dependent monooxygenase from Mycobacterium tuberculosis involved in cholesterol catabolism. J. Biol. Chem. 285, 22264–22275.

Fujii, K., Kikuchi, S., Satomi, M., Ushio-Sata, N., Morita, N., 2002. Degradation of 17β-Estradiol by a Gram-Negative Bacterium Isolated from Activated Sludge in a Sewage Treatment Plant in Tokyo, Japan. Appl. Environ. Microbiol. 68, 2057–2060.

Griffith, D.R., Kido Soule, M.C., Eglinton, T.I., Kujawinski, E.B., Gschwend, P.M., 2016. Steroidal estrogen sources in a sewage-impacted coastal ocean. Environ. Sci.: Processes Impacts. 18, 981–991.

Hamid, H., Eskicioglu, C., 2012. Fate of estrogenic hormones in wastewater and sludge treatment: A review of properties and analytical detection techniques in sludge matrix. Water Res. 46, 5813–5833.

Hanselman, T.A., Graetz, D.A., Wilkie, A.C., 2003. Manure-borne estrogens as potential environmental contaminants: a review. Environ. Sci. Technol. 37, 5471–5478.

Harvey, R.A. Ferrier, D.R., 2011. Biochemistry. Lippincott Williams & Wilkins. Baltimore, MD.

Holert, J., Cardenas, E., Bergstrand, L.H., Zaikova, E., Hahn, A.S., Hallam, S.J., Mohn, W.W., 2018. Metagenomes reveal global distribution of bacterial steroid catabolism in natural, engineered, and host environments. mBio. 9, e02345–17.

Huang, C.-H., Sedlak, D.L., 2001. Analysis of estrogenic hormones in municipal wastewater effluent and surface water using enzyme-linked immunosorbent assay and gas chromatography/tandem mass spectrometry. Environ. Toxicol. Chem. 20, 133–9.

Jurgens, M.D., Holthaus, K.I.E., Johnson, A.C., Smith, J.J.L., Hetheridge, M., Williams, R.J. 2002. The potential for estradiol and ethinylestradiol degradation in English Rivers. Environ. Toxicol. Chem. 21, 480–488.

Kao, Y.-H., Wang, S.-W., Maji, S.K., Liu, C.-W., Wang, P.-L., Chang, F.-J., Liao, C.-M., 2013. Hydrochemical, mineralogical and isotopic investigation of arsenic distribution and mobilization in the Guandu wetland of Taiwan. J. Hydrol. 498, 274–286.

Ke, J., Zhuang, W., Gin, K.Y.-H., Reinhard, M., Hoon, L.T., Tay, J.-H., 2007. Characterization of estrogen-degrading bacteria isolated from an artificial sandy aquifer with ultrafiltered secondary effluent as the medium. Appl. Microbiol. Biotechnol. 75, 1163–1171.

Kolodziej, E.P., Gray, J.L., Sedlak, D.L., 2003. Quantification of steroid hormones with pheromonal properties in municipal wastewater effluent. Environ. Toxicol. Chem. 22, 2622–2629.

Kolodziej, E.P., Harter, T., Sedlak, D.L., 2004. Dairy wastewater, aquaculture, and spawning fish as sources of steroid hormones in the aquatic environment. Environ. Sci. Technol. 38, 6377–6384.

Kramer, V.J., Miles-Richardson S., Pierens S.L., Giesy J.P., 1998. Reproductive impairment and induction of alkaline-labile phosphate, a biomarker of estrogen exposure, in fathead minnows (Pimephales promelas) exposed to waterborne 17β-estradiol. Aquat. Toxicol. 40, 335–360.

Kurisu, F., Ogura, M., Saitoh, S., Yamazoe, A., Yagi, O., 2010. Degradation of natural estrogen and identification of the metabolites produced by soil isolates of Rhodococcus sp. and Sphingomonas sp. J. Biosci. Bioeng. 109, 576–582.

Lee, Y.C., Wang, L.M., Xue, Y.H., Ge, N.C., Yang, X.M., Chen, G.H., 2006. Natural Estrogens in the Surface Water of Shenzhen and the Sewage Discharge of Hong Kong. Hum. Ecol. Risk Assess. Int. J. 12, 301–312.

Lin, A.Y., Reinhard, M., 2005. Photodegradation of common environmental pharmaceuticals and estrogens in river water. Environ. Toxicol. Chem. 24, 1303–1309.

Lin, C.-W., Wang, P.-H., Ismail, W., Tsai, Y.-W., El Nayal, A., Yang, C.-Y., Yang, F.-C., Wang, C.-H., Chiang, Y.-R., 2015. Substrate uptake and subcellular compartmentation of anoxic cholesterol catabolism in Sterolibacterium denitrificans. J. Biol. Chem. 290, 1155–1169.

Lorenzen, A., Hendel, J.G., Conn, K.L., Bittman, S., Kwabiah, A.B., Lazarovitz, G., Massé, D., McAllister, T.A., Topp, E., 2004. Survey of hormone activities in municipal biosolids and animal manures. Environ. Toxicol. 19, 216–225.

Matsumoto, T., Osada, M., Osawa, Y., Mori, K., 1997. Gonadal Estrogen Profile and Immunohistochemical Localization of Steroidogenic Enzymes in the Oyster and Scallop during Sexual Maturation. Comp. Biochem. Physiol. B, Biochem. Mol. Biol. 118, 811–817.

Morthorst, J.E., Brande-Lavridsen, N., Korsgaard, B., Bjerregaard, P., 2014. 17β-estradiol causes abnormal development in embryos of the viviparous eelpout. Environ. Sci. Technol. 48, 14668–14676.

Noguera-Oviedo, K., Aga, D.S., 2016. Chemical and biological assessment of endocrine disrupting chemicals in a full scale dairy manure anaerobic digester with thermal pretreatment. Sci. Total Environ. 550, 827–834.

Rabus, R., Widdel, F., 1995. Anaerobic degradation of ethylbenzene and other aromatic hydrocarbons by new denitrifying bacteria. Arch. Microbiol. 163, 96–103.

Shareef, A., Angove, M.J., Wells, J.D., Johnson, B.B., 2006. Aqueous solubilities of estrone, 17β-estradiol, 17α-ethynylestradiol, and bisphenol A. J. Chem. Eng. Data 51, 879–881.

Shih, C.-J., Chen, Y.-L., Wang, C.-H., Wei, S.T.-S., Lin, I.-T., Ismail, W.A., Chiang, Y.-R., 2017. Biochemical Mechanisms and Microorganisms Involved in Anaerobic Testosterone Metabolism in Estuarine Sediments. Front. Microbiol. 8, 1520.

Takamura, Y., Nomura, G., 1988. Changes in the intracellular concentration of acetyl-CoA and malonyl-CoA in relation to the carbon and energy metabolism of Escherichia coli K12. J. Gen. Microbiol. 134, 2249–2253.

Tarrant, A.M., Blomquist, C.H., Lima, P.H., Atkinson, M.J., Atkinson, S., 2003. Metabolism of estrogens and androgens by scleractinian corals. Comp. Biochem. Physiol. B Biochem. Mol. Biol. 136, 473–485.

Thayanukul, P., Zang, K., Janhom, T., Kurisu, F., Kasuga, I., Furumai, H., 2010. Concentration-dependent response of estrone-degrading bacterial community in activated sludge analyzed by microautoradiography-fluorescence in situ hybridization. Water Res. 44, 4878–4887.

Wang, P.-H., Chen, Y.-L., Wei, S.T.-S., Wu, K., Lee, T.-H., Wu, T.-Y., Chiang, Y.-R., 2020. Retroconversion of estrogens into androgens by bacteria via a cobalamin-mediated methylation. Proc. Natl. Acad. Sci. USA. 117, 1395–1403.

Wise, A., O’Brien, K., Woodruff, T., 2011. Are oral contraceptives a significant contributor to the estrogenicity of drinking water? Environ. Sci. Technol. 45, 51–60.

Wu, K., Lee, T.-H., Chen, Y.-L., Wang, Y.-S., Wang, P.-H., Yu, C.-P., Chu, K.-H., Chiang, Y.-R., 2019. Metabolites Involved in Aerobic Degradation of the A and B Rings of Estrogen. Appl. Environ. Microbiol. 85:e02223–18.

Yoshimoto, T., Nagai, F., Fujimoto, J., Watanabe, K., Mizukoshi, H., Makino, T., Kimura, K., Saino, H., Sawada, H., Omura, H., 2004. Degradation of estrogens by Rhodococcus zopfii and Rhodococcus equi isolates from activated sludge in wastewater treatment plants. Appl. Environ. Microbiol. 70, 5283–5289.

Yu, C.-P., Deeb, R.A., Chu, K.-H., 2013. Microbial degradation of steroidal estrogens. Chemosphere 91, 1225–1235.

Yu, C.-P., Roh, H., Chu, K.-H., 2007. 17β-Estradiol-Degrading Bacteria Isolated from Activated Sludge. Environ. Sci. Technol. 41, 486–492.

Zhao, H., Tian, K., Qiu, Q., Wang, Y., Zhang, H., Ma, S., Jin, S., Huo, H., 2018. Genome Analysis of Rhodococcus Sp. DSSKP-R-001: A Highly Effective β-Estradiol-Degrading Bacterium. Int. J. Genomics. 2018, 3505428.

