## Supplemental Information for "Genetic and metabolite biomarkers reveal actinobacteria-mediated estrogen biodegradation in urban estuarine sediment"

**This PDF file includes:**

Materials and Methods S1 to S3
Reference
Figures S1 to S8
Tables S1 to S3
Appendix S1 to S4

### Supplemental Materials and Methods

#### S1. Chemicals.

E1, E2, E3, 4-hydroxyestrone, 17α-ethynylestradiol, and 3a*α*-*H*-4*α*(3'-propanoate)-7a*β*-methylhexahydro-1,5-indanedione [HIP, also known as 3-(7a-methyl-1,5-dioxooctahydro-1H-inden-4-yl) propanoic acid] were purchased from Sigma-Aldrich (St. Louis, Missouri, USA). Pyridinestrone acid (PEA) was prepared as described in Chen *et al*. (Chen et al., 2017). [3,4C-$^{13}$C]E1 (99%) was purchased from Cambridge Isotope Laboratories (Tewksbury, Massachusetts, USA). All other chemicals were of analytical grade and were purchased from Honeywell Fluka (Loughborough, UK), Mallinckrodt Baker (Phillipsburg, NJ, USA), Merck (Darmstadt, Germany), and Sigma-Aldrich (St. Louis, MO, United States).

#### S2. Chemical analytical methods

##### S2.1. Ultra-performance liquid chromatography–atmospheric pressure chemical ionization–high resolution mass spectrometry (UPLC–APCI–HRMS)

Ethyl acetate extractable samples were analyzed on a UPLC system coupled to an APCI–mass spectrometer. Separation was achieved on a reversed-phase $C_{18}$ column (Acquity UPLC® BEH $C_{18}$; 1.7 µm; 100 × 2.1 mm; Waters, Milford, Massachusetts, USA) with a flow rate of 0.4 mL/min at 50℃ (column oven temperature). The mobile phase comprised a mixture of two solvents: Solvent A [2% (vol/vol) acetonitrile containing 0.1% (vol/vol) formic acid] and Solvent B [methanol containing 0.1% (vol/vol) formic acid]. Separation was achieved using a linear gradient of Solvent B from 5% to 99% across 12 min. Mass spectrum data were obtained using a Thermo Fisher Scientific™ Orbitrap

Elite$^{TM}$ Hybrid Ion Trap-Orbitrap Mass Spectrometer (Waltham, MA, USA) equipped with a standard APCI source operating in the positive ion mode. In APCI–HRMS analysis, the capillary and APCI vaporizer temperatures were 120℃ and 400℃, respectively; the sheath, auxiliary, and sweep gas flow rates were 40, 5, and 2 arbitrary units, respectively. The source voltage was 6 kV and the current was 15 μA. The parent scan was in the range of *m/z* 50–600. The predicted elemental composition of individual intermediates was calculated using Xcalibur™ Software Mass Spectrometry Software (Thermo Fisher Scientific; Waltham, MA, USA).

##### S2.2. Thin-layer chromatography (TLC)

The estrogen and estrogen metabolites were separated on silica gel aluminum TLC plates (Silica gel 60 $F_{254}$: thickness, 0.2 mm; 20 x 20 cm; Merck, Darmstadt, Hesse, Germany). Ethyl acetate: $H_2O$: acetic acid (85: 10: 10, vol/ vol/ vol) was used as the developing solvent system for the separation of PEA and HIP. Dichloromethane: ethyl acetate: ethanol (7: 2: 0.025, vol/ vol/ vol) was used as the developing solvent system for the separation of neutral estrogen metabolites. The steroid compounds were visualized under UV light at 254 nm and 302 nm or by spraying the TLC plates with 30% (vol/ vol) $H_2SO_4$.

##### S2.3. High-performance liquid chromatography (HPLC)

A reversed-phase HPLC system (Hitachi, Tokyo, Japan) was also used for identification and quantifying estrogen metabolites. Separation was achieved on an analytical RP-$C_{18}$ column (Luna $C_{18}$(2), 5 μm, 150 x 4.6 mm; Phenomenex, Torrance, California, USA) with a flow rate of 0.5 mL/min. The separation was performed isocratically at 35℃, with 40% (vol/vol) methanol containing trifluoroacetic acid (0.1%; vol/vol) serving as an eluent. The steroid products were detected in the range of 200–450 nm using a photodiode array detector.

#### S3. General molecular biological manipulations

##### S3.1. Polymerase chain reaction (PCR)

The bacterial genomic DNA (gDNA) was extracted using the Presto™ Mini gDNA Bacteria Kit (Geneaid; New Taipei City, Taiwan). PCR mixtures (25 μL) contained nuclease-free $H_2O$, 2× PCR buffer (12.5 μL), dNTPs (0.4 mM), *Taq* polymerase (5 U), Red dye and PCR stabilizer (TOOLS, Taipei, Taiwan), forward and reverse primers (each 200 nM), and template DNA (30 ng). The PCR products were verified using TAE-agarose gel (1%) electrophoresis with the SYBR® Green I nucleic acid gel stain (Thermo Fisher Scientific), and the PCR products were purified using the GenepHlow Gel/PCR Kit (Geneaid).

##### S3.2. DNA extraction and PacBio sequencing

Genomic DNA of strain B50 was obtained by using the Presto™ Mini gDNA Bacteria Kit (Geneaid; New Taipei City, Taiwan) and following the instructions. The quantity and quality of the purified DNA was evaluated by using the Nanodrop and Qubit system. The integrity of genomic DNA was examined with pulsed-field gel electrophoresis through the Pippin pulse system (Sage Science; Beverly, MA, USA). The qualified DNA was used for library construction and genome sequencing. Library construction was performed according to the protocol of SMRTbell Express template Prep Kit 2.0 (PacBio; Menlo Park, CA, USA). The DNA library was loaded onto the instrument with the Sequel

Sequencing Kit 3.0 and a SMRT Cell 1M vc3 Tray (PacBio, Menlo Park, CA, USA), and was sequenced on the PacBio Sequel platform. Raw reads were available *via* on-instrument analysis; later, post-filter subreads were obtained *via* the software, SMRT Link V8.0. These subreads were converted to HiFi reads (3 passes & 0.99 predicted accuracy) by pbccs v4.2.0. Assembly work was completed by hifiasm v0.11 and was scaffolded and polished *via* SSPACE-Long (Boetzer and Pirovano, 2014) and Arrow (https://github.com/PacificBiosciences/GenomicConsensus). When the *de novo* assembled genome was acquired, gene prediction and annotation were implemented using Prokka v1.13 (https://github.com/tseemann/prokka).

#### Supplementary Reference


Boetzer, M., Pirovano, W., 2014. SSPACE-LongRead: scaffolding bacterial draft genomes using long read sequence information. BMC Bioinformatics. 15, 211.

#### Supplemental Figures

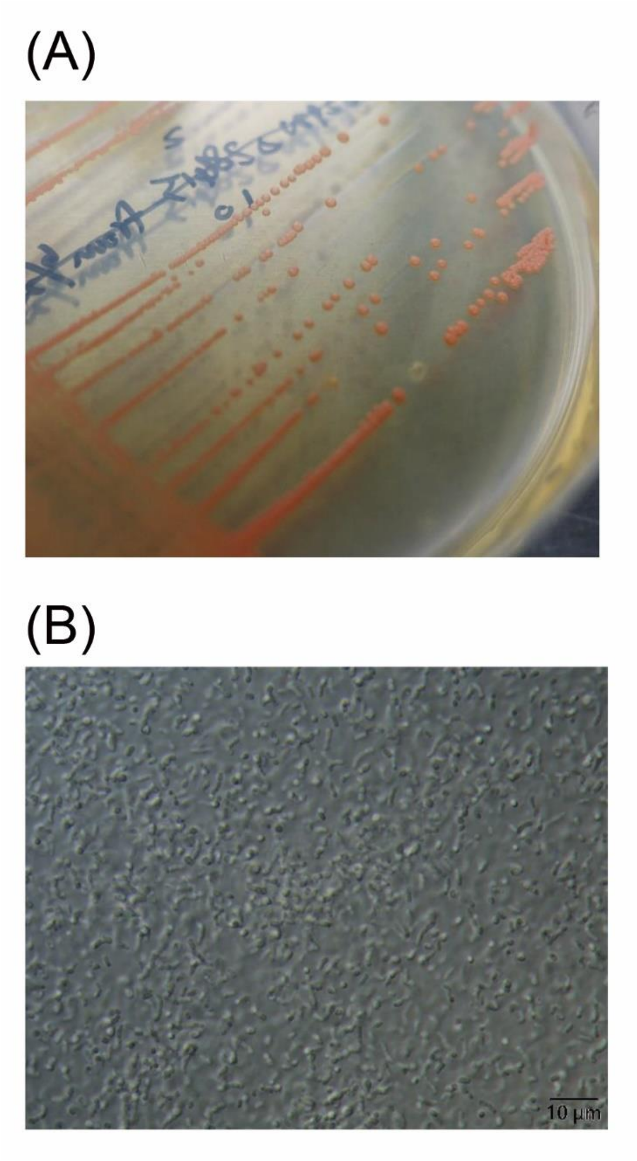


**Fig. S1.** Morphology of *Rhodococcus* sp. strain B50 colonies and cells. (A) Photograph of strain B50 colonies grown on Luria-Bertani (LB) agar. The grown colonies are smooth and contain orange pigments. (B) Light micrograph of strain B50 cells (1,000 x). Scalebar is 10 µm.

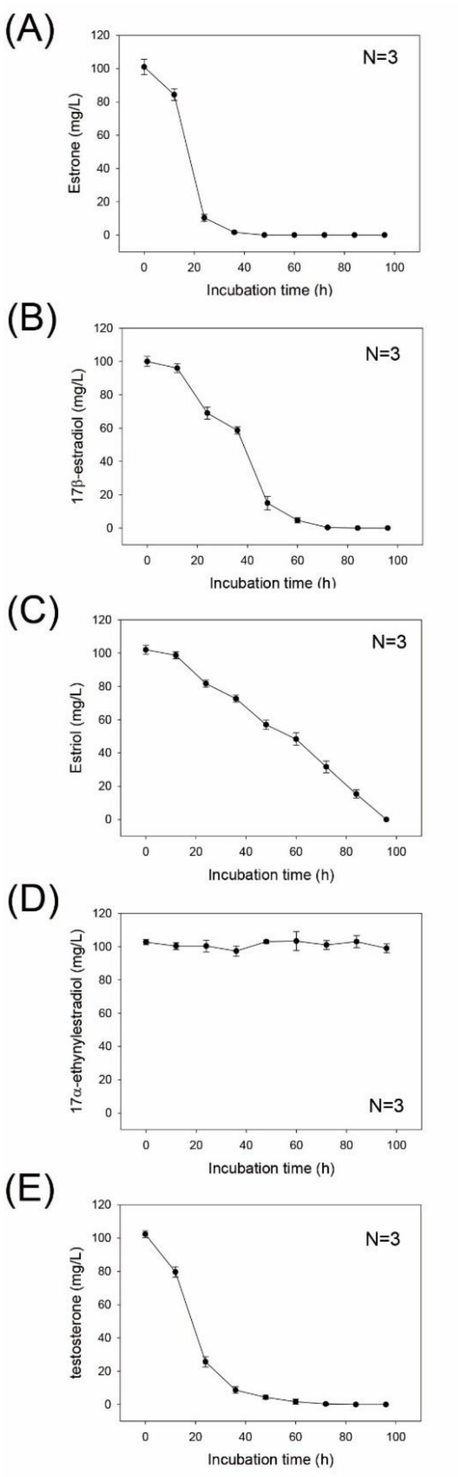


**Fig. S2.** Growth of strain B50 on several common sex steroids: (A) estrone, (B)17β-estradiol, (C) estriol, (D) 17α-ethynylestradiol, and (E) testosterone. The strain B50 cultures (50 mL) were aerobically incubated with different sex steroids in a chemically defined mineral medium. In all of the treatments, individual steroids (100 mg/L) served as the sole carbon source and electron donor. The data shown are mean values with standard errors of three experimental measurements.

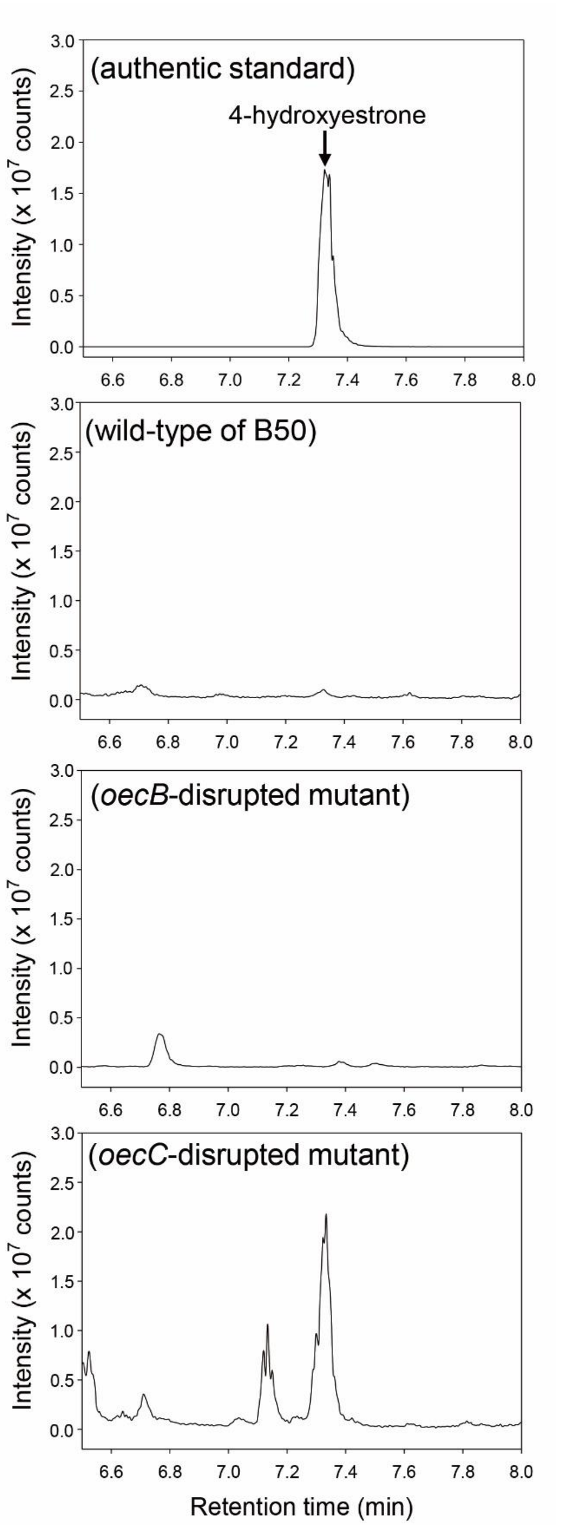


**Fig. S3.** Validation of the phenotype of the gene-disrupted strain B50 mutants. Liquid chromatography (LC) analysis of (A) an authentic 4-hydroxyestrone standard (B) the metabolite profile of the wild-type B50 strain, (C) the metabolite profile of the *oecB*-disrupted mutant, and (D) the metabolite profile of the *oecC*-disrupted mutant.

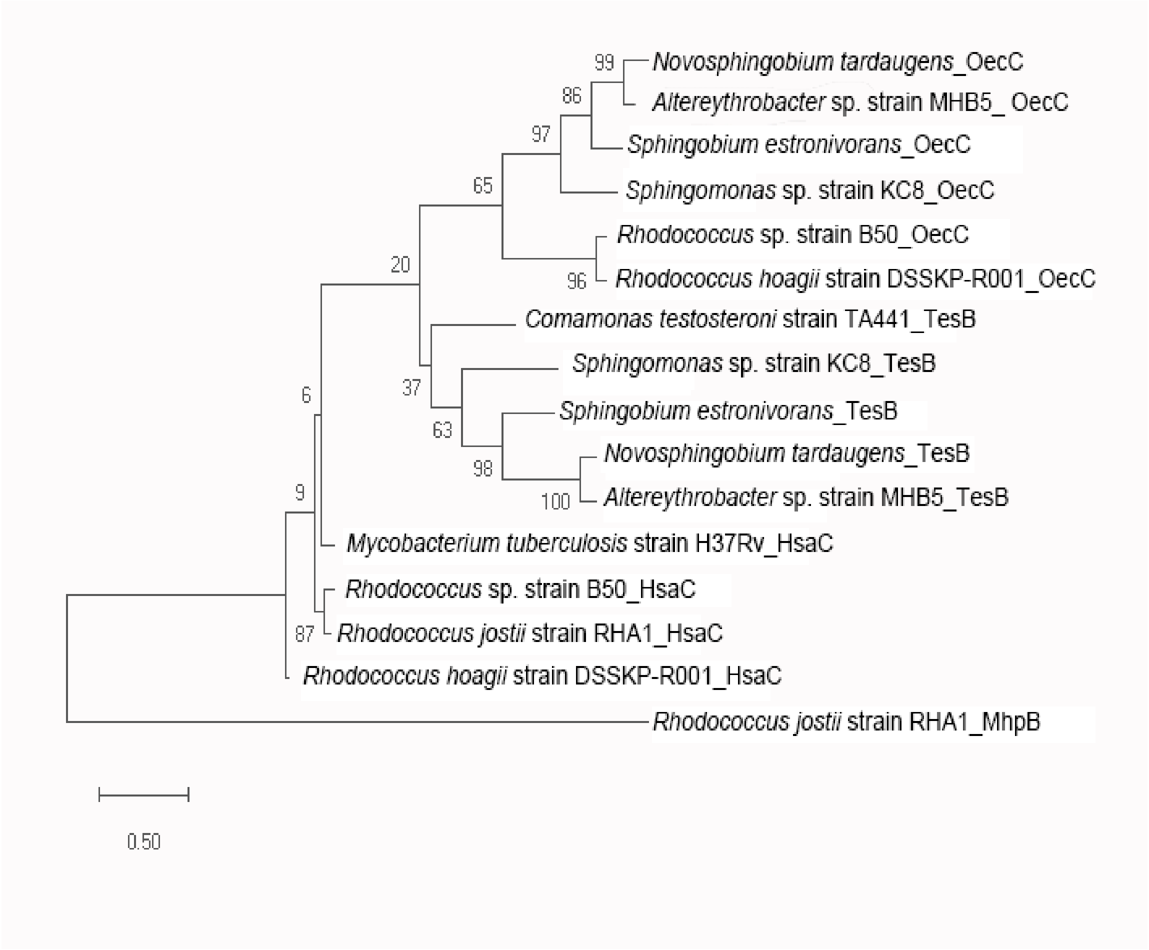


**Fig. S4.** Phylogenetic analysis of the dioxygenases involved in the *meta*-cleavage of the steroidal A-ring, including 4-hydroxyestrone 4,5-dioxygenase (OecC; estrogenic A-ring), proteobacterial TesB (androgenic A-ring), and actinobacterial HsaC (androgenic A-ring).

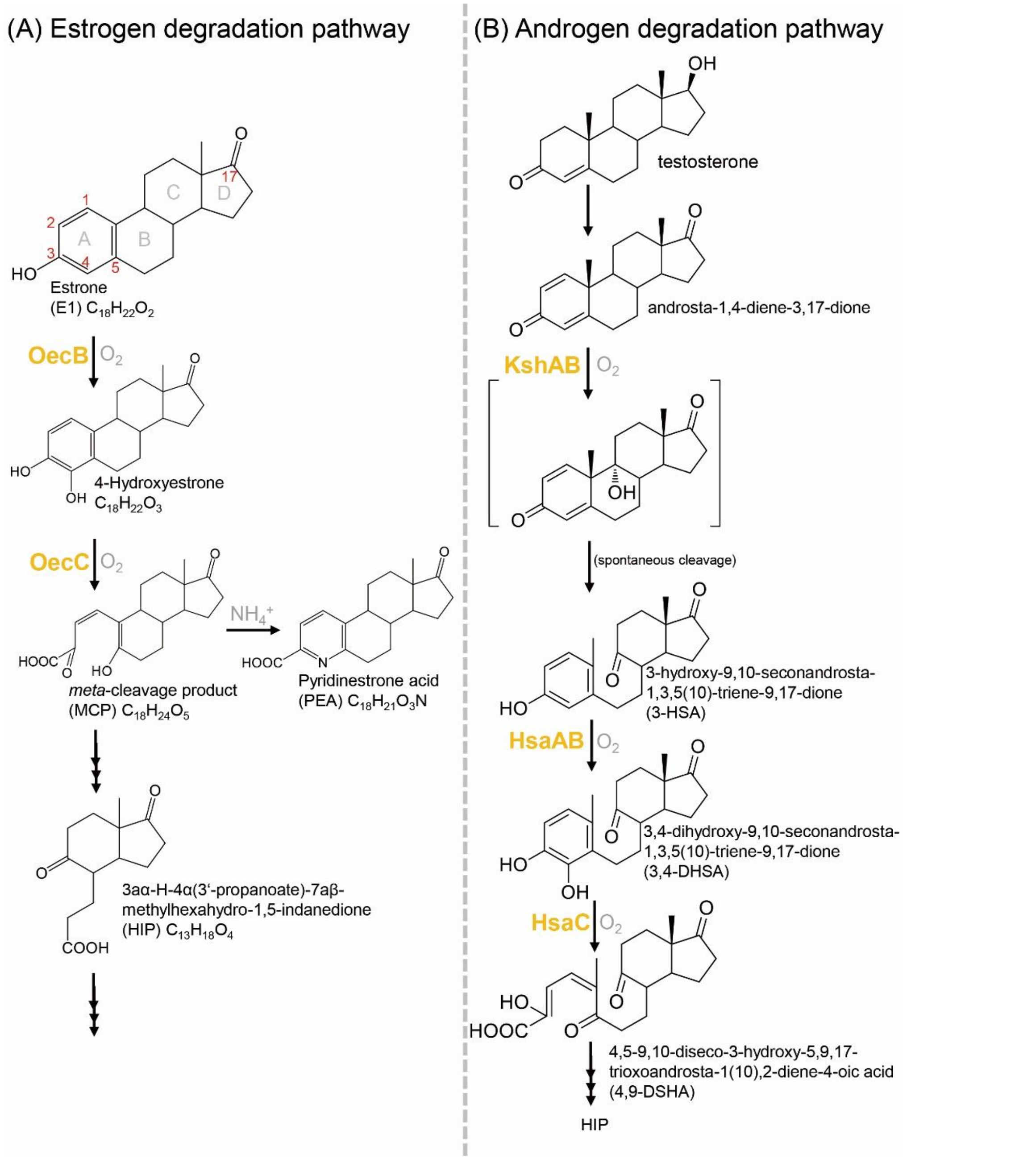


**Fig. S5.** A comparison of estrogen (A) and androgen (B) degradation pathways in actinobacteria. Characterized oxygenases are marked in orange. OecB estrone 4-hydroxylase; OecC, 4-hydroxyestrone 4,5-dioxygenase; KshAB, 3-ketosteroid 9α-hydroxylase; HsaAB, 3-hydroxy-9,10-seconandrost-1,3,5(10)-triene-9,17-dione 4-hydroxylase; HsaC, 3,4-Dihydroxy-9,10-secoandrosta-1,3,5(10)-triene-9,17-dione 4,5-dioxygenase. Protein nomenclature is based on that of *Rhodococcus* sp. strain B50 (estrogen degradation) and *R. jostii* strain RHA1 (androgen degradation). The proposed structure in the bracket (9α-hydroxy-androsta-1,4-diene-3,17-dione) is very unstable and has never been detected.

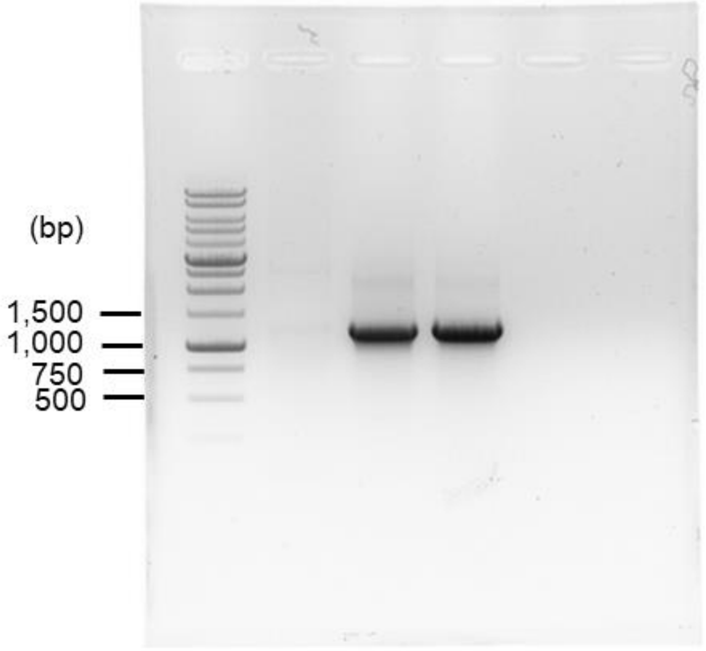


**Fig. S6.** The uncropped full-range agarose gel (1%) of Fig. 3B(i).

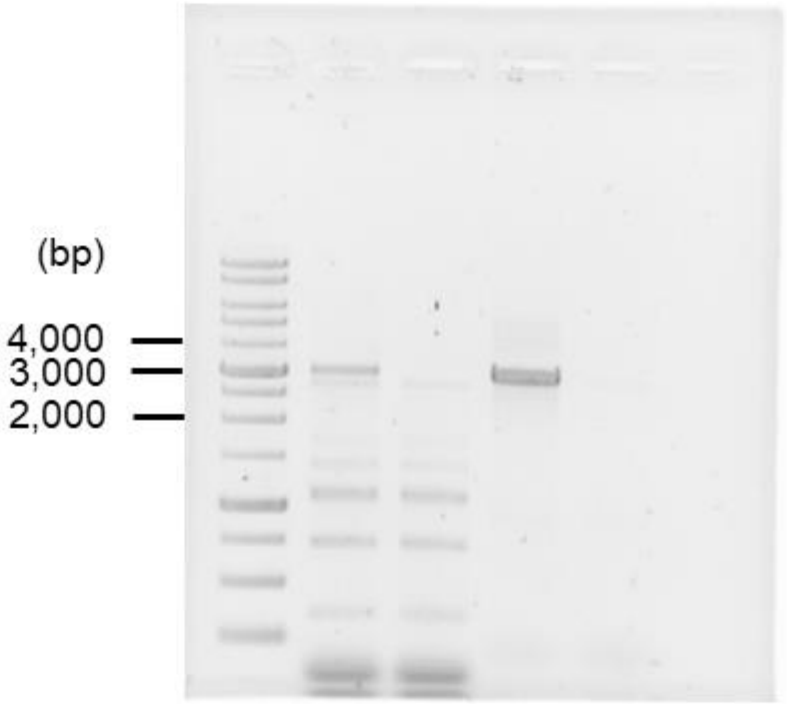


**Fig. S7.** The uncropped full-range agarose gel (1%) of Fig. 3B(ii).

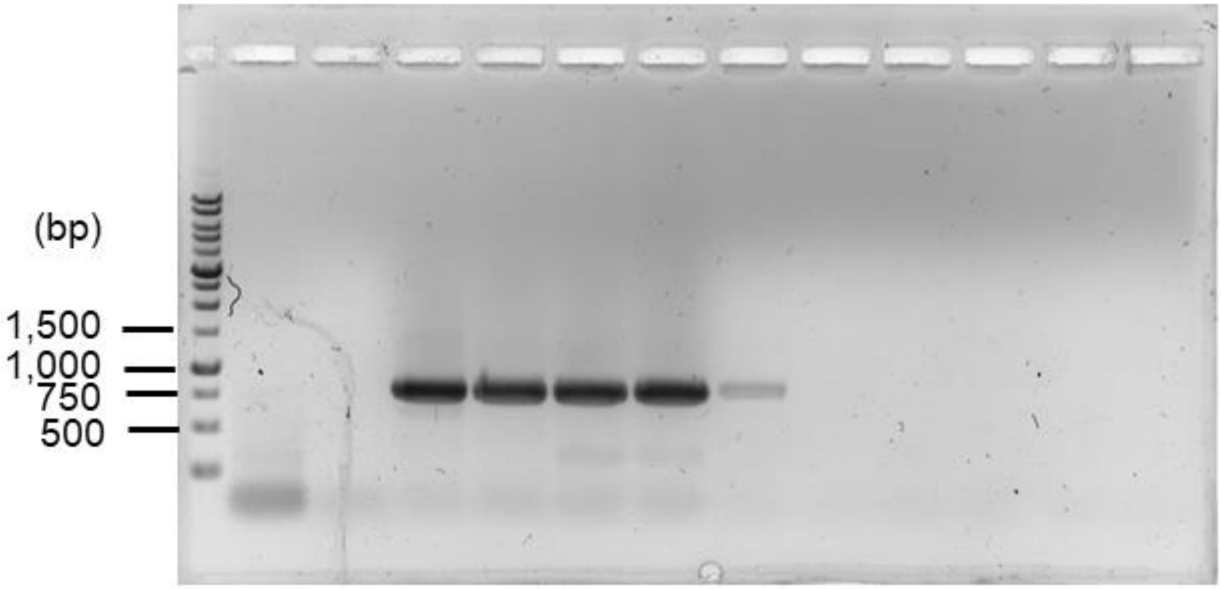


**Fig. S8.** The uncropped full-range agarose gel (1%) of Fig. 4B.

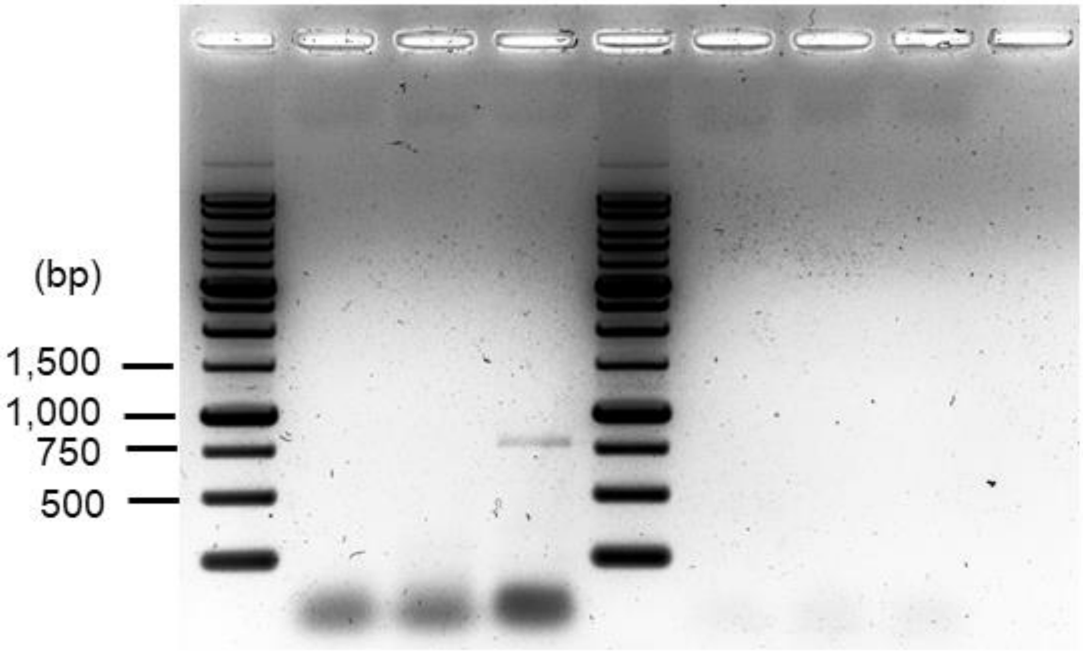


**Fig. S9.** The uncropped full-range agarose gel (1%) of Fig. 6A.

#### Supplemental Tables

**Table S1.** Antibiotic test of the *Rhodococcus* spp. reported in this study.

| Antibiotics \ Strains | B50 | A8 | 1-2-1 | 3-3-1 |
|---|---|---|---|---|
| Ampicillin (50 µg/mL)* | S | S | S | S |
| Kanamycin (25 µg/mL)* | S | R | R | R |
| Chloramphenicol (25 µg/mL)* | S | S | S | S |
| Nalidixic acid (12.5 µg/mL)* | R | R | R | R |

S, sensitive; R, resistance
*The final concentration in the growth medium of *Rhodococcus* sp.

**Table S2.** MS analysis of detected E1 metabolites in strain B50 cultures.

| Compound ID | UPLC behavior (RT[a], min) | Molecular formula/ (predicted molecular mass)[b] | Dominant ion peaks | Identification of adduct |
|---|---|---|---|---|
| E1 | 8.11 | $C_{18}H_{22}O_2$<br>270.16 | 253.16<br>271.17 | $[M-H_2O+H]^+$<br>$[M+H]^+$ |
| 4-hydroxyestrone | 7.38 | $C_{18}H_{22}O_3$<br>286.16 | 269.16<br>287.15 | $[M-H_2O+H]^+$<br>$[M+H]^+$ |
| *meta*-cleavage product | 5.27 | $C_{18}H_{24}O_5$<br>320.17 | 303.16<br>321.17 | $[M-H_2O+H]^+$<br>$[M+H]^+$ |
| pyridinestrone acid | 4.02 | $C_{18}H_{21}O_3N$<br>299.15 | 300.16 | $[M+H]^+$ |
| HIP | 3.78 | $C_{13}H_{18}O_4$<br>238.12 | 221.12<br>239.13 | $[M-H_2O+H]^+$<br>$[M+H]^+$ |

[a]RT, retention time. [b]The predicted molecular mass was calculated using the atomic mass of $^{12}C$ (12.00), $^{16}O$ (15.99), and $^{1}H$ (1.01).

**Table S3.** Oligonucleotides used in this study.

| Primer | Sequence (5'- 3') | Usage |
|---|---|---|
| oecB-UP-F<br>oecB-UP-R | ACACAGGAAACAGCTATGACCGTCGGATCGGCGCCCCG<br>AAAAGCTCCGCGATCGATACGTACGAAGTCG | For strain B50 *oecB* up-stream fragment cloning |
| oecB-DOWN-F<br>oecB-DOWN-R | GTATCGATCGCGGAGCTTTTCGTAGATCGGG<br>GTTGTAAAACGACGGCCAGTGAAGCCGAGGTCGACGCC | For strain B50 *oecB* down-stream fragment cloning |
| oecC-UP-F<br>oecC-UP-R | ACACAGGAAACAGCTATGACTTGCAATACCCGCTCGGATCG<br>GACTTCGAAGGGGCGACCTGCTCCAATCC | For strain B50 *oecC* up-stream fragment cloning |
| oecC-DOWN-F<br>oecC-DOWN-R | CAGGTCGCCCCTTCGAAGTCGAGATCGGCTGC<br>GTTGTAAAACGACGGCCAGTATGATCGCTTCCACACACGCC | For strain B50 *oecC* down-stream fragment cloning |
| Pk18-M13-F<br>Pk18-M13-R | ACTGGCCGTCGTTTTACAAC<br>GTCATAGCTGTTTCCTGTGTG | For generating the fragment of pK18-Cm$^R$-pheS** as the in-fusion backbone |
| oecB-3k-F<br>oecB-3k-R | AGGTCGATGTCCTCGACACCGAGG<br>CGCATCCTCAGTCACCTCGGCG | For checking the disruption of strain B50 *oecB* |
| oecC-3k-F<br>oecC-3k-R | AACCATGATCTTCACCATCG<br>TCAGTAGCCGTGCACGAG | For checking the disruption of strain B50 *oecC* |
| Pk18-cmpheS-F<br>Pk18-cmpheS-R | TTCATCATGCCGTTTGTGAT<br>ATCGTCAGACCCTTGTCCAC | For checking the insertion of pK18-Cm$^R$-pheS** plasmid |
| oecC-Ac-F<br>oecC-Ac-R | CGYGGCATCGGATACATCGG<br>ACMGGGTCGCAKCCGATCTC | *OecC*-specific degenerate primers for actinobacteria |
| oecC-Pr-F<br>oecC-Pr-R | CDGYYTGGGCTATSTSGG<br>ATCGCGYCSCASCCRATYTC | *OecC*-specific degenerate primers for proteobacteria |

#### Legends for Dataset

**Dataset S1 (in a separated spreadsheet).** Genome annotation of strain B50.

#### Supplemental Appendix

**Appendix S1.** Nucleotide sequence (5'-3' direction) of the 16S rRNA gene of strain B50.

ACGGAGAGTTTGATCCTGGCTCAGGACGAACGCTGGCGGCGTGCTTAACACA
TGCAAGTCGAACGATGAAGCCCAGCTTGCTGGGTGGATTAGTGGCGAACGGG
TGAGTAACACGTGGGTGATCTGCCCTGCACTTCGGGATAAGCCTGGGAAACTG
GGTCTAATACCGGATATGACCTCTTGCTGCATGGTGAGGGGTGGAAAGTTTTTC
GGTGCAGGATGAGCCCGCGGCCTATCAGCTTGTTGGTGGGGTAATGGCCTACC
AAGGCGACGACGGGTAGCCGGCCTGAGAGGGCGACCGGCCACACTGGGACT
GAGACACGGCCCAGACTCCTACGGGAGGCAGCAGTGGGGAATATTGCACAAT
GGGCGCAAGCCTGATGCAGCGACGCCGCGTGAGGGATGACGGCCTTCGGGTT
GTAAACCTCTTTCAGCAGGGACGAAGCGCAAGTGACGGTACCTGCAGAAGAA
GCACCGGCCAACTACGTGCCAGCAGCCGCGGTAATACGTAGGGTGCGAGCGT
TGTCCGGAATTACTGGGCGTAAAGAGCTCGTAGGCGGTTTGTCGCGTCGTCTG
TGAAAACCCGCAGCTCAACTGCGGGCTTGCAGGCGATACGGGCAGACTCGAG
TACTGCAGGGGAGACTGGAATTCCTGGTGTAGCGGTGAAATGCGCAGATATCA
GGAGGAACACCGGTGGCGAAGGCGGGTCTCTGGGCAGTAACTGACGCTGAG
GAGCGAAAGCGTGGGTAGCGAACAGGATTAGATACCCTGGTAGTCCACGCCG
TAAACGGTGGGCGCTAGGTGTGGGTTTCCTTCCACGGGATCCGTGCCGTAGCC
AACGCATTAAGCGCCCCGCCTGGGGAGTACGGCCGCAAGGCTAAAACTCAAA
GGAATTGACGGGGGCCCGCACAAGCGGCGGAGCATGTGGATTAATTCGATGC
AACGCGAAGAACCTTACCTGGGTTTGACATGTACCGGACGACCGCAGAGATG
TGGTTTCCCTTGTGGCCGGTAGACAGGTGGTGCATGGCTGTCGTCAGCTCGTG
TCGTGAGATGTTGGGTTAAGTCCCGCAACGAGCGCAACCCTTGTCCTGTGTTG
CCAGCACGTGATGGTGGGGACTCGCAGGAGACTGCCGGGGTCAACTCGGAGG
AAGGTGGGGACGACGTCAAGTCATCATGCCCCTTATGTCCAGGGCTTCACACA
TGCTACAATGGTCGGTACAGAGGGCTGCGATACCGTGAGGTGGAGCGAATCCC
TTAAAGCCGGTCTCAGTTCGGATCGGGGTCTGCAACTCGACCCCGTGAAGTCG
GAGTCGCTAGTAATCGCAGATCAGCAACGCTGCGGTGAATACGTTCCCGGGCC
TTGTACACACCGCCCGTCACGTCATGAAAGTCGGTAACACCCGAAGCCGGTG
GCCTAACCCCTCGTGGGAGGGAGCCGTCGAAGGTGGGATCGGCGATTGGGAC
GAAGTCGTAACAAGGTAGCCGTACCGGAAGGTGCGGCTGGATCACCTCCTTT

**Appendix S2.** Nucleotide sequence (5'-3' direction) of the *oecB* gene (GMFMDNLD_3_05336) of strain B50.

ATGACTTCCGATATTTCGGCAACCGATCAGGCAGCTGTCGACTACGATCCGTTC
GCACCCGAAGCCATGGTCGACCCTCGCCCGATCTACGAAAAGCTCCGTGCTGC
AGGTCCTTTGCACTACCTACCCCAGTACGATGCGTGGGCGTTGTGCAGCTTCG
AAGCCGTTTGGCGCGTCACCCGGGATCTGAAGAACTTCACGACCGAGCATCG
CGGTTCGCCGCCGATGAATTCGTTGCTCGGCGAACCCAGTTTCCCCAATTTCA
CCCAGACCGATCCGCGCGAGCACGCTGCTGCTCGCAGGCTGCTCCAACCCGC
CTACAACAAGGCGGCGGCGGATCACGACGCGGCATACATGCGTACGCTCGCTC
GTGAAGTCATCACACCGCTCGTCGAGGGTGGCGACGGCACCATGGATGTATTC
CGCGATTACGCCAGCCGTGTCGCCGCTCGCTTCGCCGGCCACAAGGCCGGCCT
TCCCGCCGCCGACTCGGAGCGGATTCGGCACCGTGCCGAACAACTCTTCGTCC
GCGAATACGGGCAGCGTGGCACGTCCCCGAGCAATGCCGAAGCAGGTGCCGA
AGTATTCGAATACCTTCAAGAGCTGGTCGTCGAGGCTCGTCGGGACCCGTCGA
GCGCCCGAGGTGATCTGGCGTCCTTGCTCAACGGCACCGTCCGGGGTCGTTCT
CTGACCGATCAGGAAATCCTCGGCAACTTGCTCACCCTGATCATCACCGGCTC
CGAAACGACCGAGATCAGCGTCGCAGCGACCCTGTACTACCTCGCCCGAAAT
CCTGAGCAGCTTGCCGCGGTTCGCGCCGACCGCAGCCTGCTTCTCAATGCCTT
CATGGAAACCGTGCGATTCGACCACCCGACCGACATTCTATGTCGGGAGGTGG
TCAACGAGGTCGAGGTCTGCGGCCGAAAGCTCCTCCCCGGCCAACAACTCAT
CCTCATGTGGGGTTCTGCAGGTCGCGATGAAAGCGAGTTCCCGGACGCCGAC
ACCTACGATATCCACCGCACCTACCAGCGGCACCTGCTCTTCGGGCACGGCCA
GCACAAGTGCCTCGGCGAGAGCATCGCACTTCGGCTCGGCACGATCATGCTCG
AAGAGTTCTTCGAGGCGATCGATACGTACGAAGTCGACTGGGACGGCTGCCG
CCGCAAATACGCCGAATTCGTCCAGGGGTTCAACTCCGTCCCCATCCGATTCA
CCACCGCCTGA

**Appendix S3.** Nucleotide sequence (5'-3' direction) of the *oecC* gene (GMFMDNLD_3_05338) of strain B50.

ATGAATCTTCGCGGCATCGGATACATCGGACTCGACGTCCCCAACCGACAGGA
ATGGGACCGATTCGCCACCACCGTCGTGGGATTGGAGCAGGTCGCCCGTCCG
GCCGGGGATGACGACGCGTCCTACTTCAAGGCCGATGATCGCAGCTGGCGTAT
CGCGGCCCGAGAGAGCAACACCCCCGGCATCGGATTCATCGGATTCGAAGTC
AACGACCGCGCCAGCTTCGACGACGCGGTCCGCATCCTCGAGACAGCCGGCG
CCGCACCCAAGGCTGCCGCCGAAACCGAACTCGCCGAACGAGGCGTGCAGG
CCATGGTCTGGTTCGAAGATCCTGCAGGGACCCGACTGGAGATCTTCTGGGGC
CCGACCGTCGACGGCGCATTCCGCTCTCCGCTGGGCGAGCCGGGCTTCGTCAC
TGAAGGTGGATTCGGCCACGCGGTGTTGATGGTGCAGGATCTGCCCGCCGCCC
TCGAGTTCTACACGTCCGTGCTGGGCATGCGCACCTCCGACTTCATGAACTTC
GGCGAGGGCATGGCCATTCACTTCCTGCGATGCACCCCGCGCCATCACAGCAT
CGCGCTCAGCGCCGTCGGTCCGGTCTCGGGAACCCACCACATCGCGCTCGAG
GTCGCCGATGTCGATCAGGTGGGCACAGCACTCGACCGCGCAACCGAAGCGG
GTCTGGCCATCACCGCATCGCTCGGCCGCCACAAGAACGACCGCATGCTCTCG
TTCTACATGCGCAGCCCCGCGGGCTTCGAAGTCGAGATCGGCTGCGACCCGGT
CCTCGTCGACGAGGAAACGTGGATCACCAACGAGTTCACCGGCGGTGACGCC
TGGGGGCACCACGGACTGACCAGCGAATCACTCGCCGAGTCGGTCGCCGGGA
ACGCATCATGA

**Appendix S4.** Nucleotide sequences (5'-3' direction) of the *oecC* amplicons identified in the E1-spiked sediment.

>sediment_cDNA#1
CGCGGCATCGGATACATCGGCCTGAACGTCCCGGACCGGCAGGAATGGGACA
AGTTCGCCACCGACGTCATCGGACTCCAGACGGTTACCCTCCCGGCCGGCGCC
GACGACGCGTCCTACTTCAAGGCCGACACTCGAAGCTGGCGTCTCGCCGTGC
GCGAGAACTCCATTCCCGGAGTCGATTTCATCGGTTTCGAGGTACTGGACGTC
CACGGCTTCGACGAGGCGGTCCAGGAGCTCGAGGCAGCAGGTGCAGCGCCG
AAGCGCGCCAGCGACTGGGAACTGGCCGAGCGCGGCGTCCAGGCGATGGTGT
GGTTCGAGGATCCCGCGGGCACCCGGCTCGAGATCTTCTGGGGCCCCACCGTC
GACGGCGCCTTCCGCTCGCCGGCCGGCGTCCCGGGATTCGTGACCGAGGGCG
GATTCGGGCACGCGGTGATGATGGTGCTCGATCTCCCCGCCGCGCTGGAGTTC
TACTCCACGGTGCTGGGGATGCGCACCTCGGACTTCATGAACTTCGGCGAGGG
CATGGCGATCCACTTCCTGCGCTGTACGCCGCGCCACCACACCATCGCACTCA
GCGCCGTGGGACCGATCTCGGGCACCCACCACATCGCCTTCGAAGTGGCCGA
GGTCGACCAGGTCGGCGCCGCCCTCGACCGCGCAACCCAGGCAGGCCTCGCC
GTCACCGCGTCACTCGGCCGGCACAAGAACGACCGCATGCTCTCCTTCTACAT
GCGTAGCCCCGCGGGTTTCGAGGTCGAGATCGGCTGCGACCCTGT
>sediment_cDNA#2
GGATACATCGGCCTGAACGTCCCGGACCGGCAGGAATGGGACAAGTTCGCCA
CCGACGTCATCGGACTCCAGACGGTTACCCTCCCGGCCGGCGCCGACGACGC
GTCCTACTTCAAGGCCGACACTCGAAGCTGGCGTCTCGCCGTGCGCGAGAAC
TCCATTCCCGGAGTCGATTTCATCGGTTTCGAGGTACTGGACGTCCACGGCTTC
GACGAGGCGGTCCAGGAGCTCGAGGCAGCAGGTGCAGCGCCGAAGCGCGCC
AGCGACCGGGAACTGGCCGAGCGCGGCGTCCAGGCGATGGTGTGGTTCGAGG
ATCCCGCGGGCACCCGGCTCGAGATCTTCTGGGGCCCCACCGTCGACGGCGC
CTTCCGCTCGCCGGCCGGCGTCCCGGGATTCGTGACCGAGGGCGGATTCGGG
CACGCGGTGATGATGGTGCTCGATCTCCCCGCCGCGCTGGAGTTCTACTCCAC
GGTGCTGGGGATGCGCACCTCGGACTTCATGAACTTCGGCGAGGGCATGGCG
ATCCACTTCCTGCGCTGTACGCCGCGCCACCACACCATCGCACTCAGCGCCGT
GGGACCGATCTCGGGCACCCACCACATCGCCTTCGAAGTGGCCGAGGTCGAC
CAGGTCGGCGCCGCCCTCGACCGCGCAACCCAGGCAGGCCTCGCCGTCACCG
CGTCACTCGGCCGGCACAAGAACGACCGCATGCTCTCCTTCTACATGCGTAGC
CCCGCGGGTTTCGAGGTCGAGATCGGCTGCGACCCGGT
>sediment_cDNA#3
TCGGACTCGACGTCCCCAACCGACAGGAATGGGACCGATTCGCCACCACCGT
CGTGGGATTGGAGCAGGTCGCCCGTCCGGCCGGGGATGACGACGCGTCCTAC
TTCAAGGCCGATGATCGCAGCTGGCGTATCGCGGCCCGAGAGAGCAACACCC
CCGGCATCGGATTCATCGGATTCGAAGTCAACGACCGCGCCAGCTTCGACGAC
GCGGTCCGCATCCTCGAGGCAGCCGGCGCCGCACCCAAGGCTGCCGCCGAAA
CCGAACTCGCCGAACGAGGCGTGCAGGCCATGGTCTGGTTCGAAGATCCTGC
AGGGACCCGACTGGAGATCTTCTGGGGCCCGACCGTCGACGGCGCATTCCGC
TCTCCGCTGGGCGAGCCGGGCTTCGTCACTGAAGGTGGATTCGGCCACGCGG
CGTTGATGGTGCAGGATCTGCCCGCCGCCCTCGAGTTCTACACGTCCGTGCTG
GGCATGCGCACCTCCGACTTCATGAACTTCGGCGAGGGCATGGCCATTCACTT

CCTGCGATGCACCCCGCGCCATCACAGCATCGCGCTCAGCGCCGTCGGTCCAG
TCTCGGGAACCCACCACATCGCGCTCGAGGTCGCCGATGTCGATCAGGTGGGC
ACAGCACTCGACCGCGCAACCGAAGCGGGTCTGGCCATCACCGCATCGCTCG
GCCGCCACAAGAACGACCGCATGCTCTCGTTCTACATGCGCAGCCCCGCGGGC
TTCGAAGTCGAGATCGGCTGCGACCCTGT
>sediment_cDNA#4
TCGCCACCACCGTCGTGGGATTGGAGCAGGTCGCCCGTCCGGCCGGGGATGA
CGACGCGTCCTACTTCAAGGCCGATGATCGCAGCTGGCGTATCGCGGCCCGAG
AGAGCAACACCCCCGGCATCGGATTCATCGGATTCGAAGTCAACGACCGCGC
CAGCTTCGACGACGCGGTCCGCATCCTCGAGACAGCCGGCGCCGCACCCAAG
GCTGCCGCCGAAACCGAACTCGCCGAACGAGGCGTGCAGGCCATGGTCTGGT
TCGAAGATCCTGCAGGGACCCGACTGGAGATCTTCTGGGGCCCGACCGTCGA
CGGCGCATTCCGCTCTCCGCTGGGCGAGCCGGGCTTCGTCACTGAAGGTGGAT
TCGGCCACGCGGCGTTGATGGTGCAGGATCTGCCCGCCGCCCTCGAGTTCTAC
ACGTCCGTGCTGGGCATGCGCACCTCCGACTTCATGAACTTCGGCGAGGGCAT
GGCCATTCACTTCCTGCGATGCACCCCGCGCCATCACAGCATCGCGCTCAGCG
CCGTCGGTCCGGTCTCGGGAACCCACCACATCGCGCTCGAGGTCGCCGATGTC
GATCAGGTGGGCACAGCACTCGACCGCGCAACCGAAGCGGGTCTGGCCATCA
CCGCATCGCTCGGCCGCCACAAGAACGACCGCATGCTCTCGTTCTACATGCGC
AGCCCCGCGGGCTTCGAGGTCGAGATCGGCTGCGACCCTGT
>sediment_cDNA#5
AATTCCAAGCTTCGCGGCATTGGATACATCGGCCTGAACGTCCCGGACCGGCA
GGAATGGGACAAGTTCGCCACCGACGTCATCGGACTCCAGACGGTTACCCTCC
CGGCCGGCGCCGACGACGCGTCCTACTTCAAGGCCGACACTCGAAGCTGGCG
TCTCGCCGTGCGCGAGAACTCCATTCCCGGAGTCGATTTCATCGGTTTCGAGG
TACTGGACGTCCACGGCTTCGACGAGGCGGTCCAGGAGCTCGAGGCAGCAGG
TGCAGCGCCGAAGCGCGCCAGCGACCGGGAACTGGCCGAGCGCGGCGTCCA
GGCGATGGTGTGGTTCGAGGATCCCGCGGGCACCCGGCTCGAGATCTTCTGGG
GCCCCACCGTCGACGGCGCCTTCCGCTCGCCGGCCGGCGTCCCGGGATTCGTG
ACCGAGGGCGGATTCGGGCACGCGGTGATGATGGTGCTCGATCTCCCCGCCGC
GCTGGAGTTCTACTCCACGGTGCTGGGGATGCGCACCTCGGACTTCATGAACT
TCGGCGAGGGCATGGCGATCCACTTCCTGCGCTGTACGCCGCGCCACCACACC
ATCGCACTCAGCGCCGTGGGACCGATCTCGGGCACCCACCACATCGCCTTCGA
AGTGGCCGAGGTCGACCAGGTCGGCGCCGCCCTCGACCGCGCAACCCAGGCA
GGCCTCGCCGTCACCGCGTCACTCGGCCGGCACAAGAACGACCGCATGCTCT
CCTTCTACATGCGTAGCCCCGCGGGTTTCGAGGTCGAGATCGGCTGCGACCCG
GT
>sediment_cDNA#6
CGTGGCATCGGATACATCGGCCTGAACGTCCCGGACCGGCAGGAATGGGACA
AGTTCGCCACCGACGTCATCGGACTCCAGACGGTTACCCTCCCGGCCGGCGCC
GACGACGCGTCCTACTTCAAGGCCGACACTCGAAGCTGGCGTCTCGCCGTGC
GCGAGAACTCCATTCCCGGAGTCGATTTCATCGGTTTCGAGGTACTGGACGTC
CACGGCCTCGACGAGGCGGTCCAGGAACTCGAGGCAGCAGGTGCGGCGCCG
AAGCGCGCCAGCGACCGGGAACTGGCCGAGCGCGGCGTCCAGGCGATGGTG
TGGTTCGAGGATCCCGCGGGCACCCGGCTCGAGATCTTCTGGGGCCCCACCGT
CGACGGCGCCTTCCGCTCGCCGGCCGGCGTCCCGGGATTCGTGACCGAGGGC

GGATTCGGGCACGCGGTGATGATGGTGCTCGATCTCCCCGCCGCGCTGGAGTT
CTACTCCACGGTGCTGGGGATGCGCACCTCGGACTTCATGAACTTCGGCGAGG
GCATGGCGATCCACTTCCTGCGCTGTACGCCGCGCCACCACACCATCGCACTC
AGCGCCGTGGGACCGATCTCGGGCACCCACCACATCGCCTTCGAAGTGGCCG
AGGTCGACCAGGTCGGCGCCGCCCTCGACCGCGCAACCCAGGCAGGCCTCGC
CGTCACCGCGTCACTCGGCCGGCACAAGAACGACCGCATGCTCTCCTTCTACA
TGCGTAGCCCCGCGGGTTTCGAGGTCGAGATCGGCTGCGACCCTGT
>sediment_cDNA#7
CGCGGCATCGGATACATCGGACACGACGTCCCCAACCGACAGGAATGGGACC
GATTCGCCACCACCGTCGTGGGATTGGAGCAGGTCGCCCGTCCGGCCGGGGA
TGACGACGCGTCCTACTTCAAGGCCGATGATCGCAGCTGGCGTATCGCGGCCC
GAGAGAGCAACACCCCCGGCATCGGATTCATCGAATTCGAAGTCAACGACCG
CGCCAGCTTCGACGACGCGGTCCGCATCCTCGAGACAGCCGGCGCCGCACCC
AAGGCTGCCGCCGAAACCGAACTCGCCGAACGAGGCGTGCAGGCCATGGTCT
GGTTCGAAGATCCTGCAGGGACCCGACTGGAGATCTTCTGGGGCCCGACCGT
CGACGGCGCATTCCGCTCTCCGCTGGGCGAGCCGGGCTTCGTCACTGAAGGT
GGATTCGGCCACGCGGCGTTGATGGTGCAGGATCTGCCCGCCGCCCTCGAGTT
CTACACGTCCGTGCTGGGCATGCGCACCTCCGACTTCATGAACTTCGGCGAGG
GCATGGCCATTCACTTCCTGCGATGCACCCCGCGCCATCACAGCATCGCGCTC
AGCGCCGTCGGTCCGGTCTCGGGAACCCACCACATCGCGCTCGAGGTCGCCG
ATGTCGATCAGGTGGGCACAGCACTCGACCGCGCAACCGAAGCGGGTCTGGC
CATCACCGCATCGCTCGGCCGCCACAAGAACGACCGCATGCTCTCGTTCTACA
TGCGCAGCCCCGCGGGCTTCGAAGTCGAGATCGGCTGCGACCCTGT
>sediment_cDNA#8
CGCGGCATCGGATACATCGGCCTGAACGTCCCGGACCGGCAGGAATGGGACA
AGTTCGCCACCGACGTCATCGGACTCCAGACGGTTACCCTCCCGGCCGGCGCC
GACGACGCGTCCTACTTCAAGGCCGACACTCGAAGCTGGCGTCTCGCCGTGC
GCGAGAACTCCATTCCCGGAGTCGATTTCATCGGTTTCGAGGTACTGGACGTC
CACGGCTTCGACGAGGCGGTCCAGGAGCTCGAGGCAGCAGGTGCAGCGCCG
AAGCGCGCCAGCGACCGGGAACTGGCCGAGCGCGGCGTCCAGGCGATGGTG
TGGTTCGAGGATCCCGCGGGCACCCGGCTCGAGATCTTCTGGGGCCCCACCGT
CGACGGCGCCTTCCGCTCGCCGGCCGGCGTCCCGGGATTCGTGACCGAGGGC
GGATTCGGGCACGCGGTGATGATGGTGCTCGATCTCCCCGCCGCGCTGGAGTT
CTACTCCACGGTGCTGGGGATGCGCACCTCGGACTTCATGAACTTCGGCGAGG
GCATGGCGATCCACTTCCTGCGCTGTACGCCGCGCCACCACACCATCGCACTC
AGCGCCGTGGGACCGATCTCGGGCACCCACCACATCGCCTTCGAAGTGGCCG
AGGTCGACCAGGTCGGCGCCGCCCTCGACCGCGCAACCCAGGCAGGCCTCGC
CGTCACCGCGTCACTCGGCCGGCACAAGAACGACCGCATGCTCTCCTTCTACA
TGCGTAGCCCCGCGGGTTTCGAGGTCGAGATCGGCTGCGACCCTGT
>sediment_cDNA#9
CGCGGCATCGGATACATCGGACTCGACGTCCCCAACCGACAGGAATGGAACC
GATTCGCCACCACCGTCGTGGGATTGGAGCAGGTCGCCCGTCCGGCCGGGGA
TGACGACGCGTCCTACTTCAAGGCCGATGATCGCAGCTGGCGTATCGCGGCCC
GAGAGAGCAACACCCCCGGCATCGGATTCATCGGATTCGAAGTCAACGACCG
CGCCAGCTTCGACGACGCGGTCCGCATCCTCGAGACAGCCGGCGCCGCACCC
AAGGCTGCCGCCGAAACCGAACTCGCCGAACGAGGCGTGCAGGCCATGGTCT

GGTTCGAAGATCCTGCAGGGACCCGACTGGAGATCTTCTGGGGCCCGACCGT
CGACGGCGCATTCCGCTCTCCGCTGGGCGAGCCGGGCTTCGTCACTGAAGGT
GGATTCGGCCACGCGGTGTTGATGGTGCAGGATCTGCCCGCCGCCCTCGAGTT
CTACACGTCCGTGCTGGGCATGCGCACCTCCGACTTCATGAACTTCGGCGAGG
GCATGGCCATTCACTTCCTGCGATGCACCCCGCGCCATCACAGCATCGCGCTC
AGCGCCGTCGGTCCGGTCTCGGGAACCCACCACATCGCGCTCGAGGTCGCCG
ATGTCGATCAGGTGGGCACAGCACTCGACCGCGCAACCGAAGCGGGTCTGGC
CATCACCGCATCGCTCGGCCGCCACAAGAACGACCGCATGCTCTCGTTCTACA
TGCGCAGCCCCGCGGGCTTCGAAGTCGAGATCGGCTGCGACCCTGT
>sediment_cDNA#10
TCCCGGACCGGCAGGAATGGGACAAGTTCGCCACCGACGTCATCGGACTCCA
GACGGTTACCCTCCCGGCSGGCGCCGACGACGCGTCCTACTTCAAGGCCGAC
ACTCGAAGCTGGCGTCTCGCCGTGCGCGAGAACTCCATTCCCGGAGTCGATTT
CACCGGTTTCGAGGTACTGGACGTCCACGGCTTCGACGAGGCGGTCCAGGAG
CTCGAGGCAGCAGGTGCAGCGCCGAAGCGCGCCAGCGACCGGGAACTGGCC
GAGCGCGGCGTCCAGGCGATGGTGTGGTTCGAGGATCCCGCGGGCACCCGGC
TCGAGACCTTCTGGGGCCCCACCGTCGACGGCGCCTTCCGCTCGCCGGCCGG
CGTCCCGGGATTCGTGACCGAGGGCGGATTCGGGCACGCGGTGATGATGGTG
CTCGATCTCCCCGCCGCGCTGGAGTTCTACTCCACGGTGCTGGGGATGCGCAC
CTCGGACTTCATGAACTTCGGCGAGGGCATGGCGATCCACTTCCTGCGCTGTA
CGCCGCGCCACCACACCATCGCACTCAGCGCCGTGGGACCGATCTCGGGCAC
CCACCACATCGCCTTCGAAGTGGCCGAGGTCGACCAGGTCGGCGCCGCCCTC
GACCGCGCAACCCAGGCAGGCCTCGCCGTCACCGCGTCACTCGGCCGGCACA
AGAACGACCGCATGCTCTCCTTCTACATGCGTAGCCCCGCGGGTTTCGAGGTC
GAGATCGGCTGCGACCCTGT
